# A large-scale Boolean model of the Rheumatoid Arthritis Fibroblast-Like Synoviocytes predicts drug synergies in the arthritic joint

**DOI:** 10.1101/2023.01.16.524300

**Authors:** Vidisha Singh, Aurelien Naldi, Sylvain Soliman, Anna Niarakis

**Affiliations:** Université Paris-Saclay, Laboratoire Européen de Recherche pour la Polyarthrite rhumatoïde - Genhotel, Univ Evry, Evry, France; Lifeware Group, Inria, Saclay-île de France, 91120 Palaiseau, France

**Keywords:** Computational systems biology, Boolean Network, Rheumatoid Arthritis Synovial Fibroblasts, *in silico* simulations, drug repurposing, theurapeutic targets, drug candidates

## Abstract

Rheumatoid arthritis (RA) is a complex autoimmune disease with an unknown aetiology. However, rheumatoid arthritis fibroblast-like synoviocytes (RA-FLS) play a significant role in initiating and perpetuating destructive joint inflammation by expressing immuno-modulating cytokines, adhesion molecules, and matrix remodelling enzymes. In addition, RA-FLS are primary drivers of inflammation, displaying high proliferative rates and an apoptosis-resistant phenotype. Thus, RA-FLS-directed therapies could become a complementary approach to immune-directed therapies by predicting the optimal conditions that would favour RA-FLS apoptosis, limit inflammation, slow the proliferation rate and minimise bone erosion and cartilage destruction. In this paper, we present a large-scale Boolean model for RA-FLS that consists of five submodels focusing on apoptosis, cell proliferation, matrix degradation, bone erosion and inflammation. The five phenotype-specific submodels can be simulated independently or as a global model. *In-silico* simulations and perturbations reproduced the expected biological behaviour of the system under defined initial conditions and input values. The model was then used to mimic the effect of mono or combined therapeutic treatments and predict novel targets and drug candidates through drug repurposing analysis.

## Introduction

Rheumatoid arthritis (RA) is a multifactorial disease that affects the articular joints of the human body and involves combinations of genetic factors and environmental triggers (McInnes & Schett, 2011, 2007; Ostrowska *et al*, 2018). Cytokines such as tumour necrosis factor-alpha (TNF-α) and interleukin 6 (IL-6) are central players in RA, causing inflammation and tissue damage, whereas their inhibitors are considered among the main treatments for RA (McInnes & Schett, 2007; Noack & Miossec, 2017; Alunno *et al*, 2017). However, despite a growing number of such drugs, 40% of patients fail to respond to therapy adequately (Wijbrandts & Tak, 2017; Rein & Mueller, 2017). Recent studies have identified RA fibroblast-like synoviocytes (RA-FLS) as responsible for up to a quarter of the disease’s heritability (Ge *et al*, 2021), attributing to these cells a causal role in disease pathogenesis. The primary roles of RA-FLS in RA are discussed below (Bartok & Firestein, 2010; Turner & Filer, 2015). RA-FLS plays a central role in the pathogenesis of RA by activating the innate immune response. The immune response is maintained via the secretion of soluble molecules, such as proinflammatory cytokines IL-6, IL-1, and TNF, in response to environmental stimuli and interactions with other cells. Secretions of these molecules act as a positive feedback loop and eventually trigger the activation of RA-FLS into expressing the responses repeatedly (Bartok & Firestein, 2010). Joint inflammation is the primary characteristic of RA. During joint inflammation in RA, FLS proliferate to form the pannus, which invades and destroys the cartilage. Major pathways involved in the perpetuation of inflammation are TNF, IL-6, IL-1, and IL-17 (Choy & Panayi, 2001; Noack & Miossec, 2017).

The final stages of the disease involve the degradation of bone and cartilage due to chronic inflammation within the joint area. During chronic and sustained inflammation, RA-FLS secrete two groups of soluble molecules: a) receptor activator of nuclear factor kappa-B ligand (RANKL), a molecule that promotes osteoblasts differentiation to osteoclasts, cells that are responsible for bone degradation and bone resorption (Boyle *et al*, 2003; Sato & Takayanagi, 2006) and b) matrix metalloproteinases (MMPs), a group of matrix proteases responsible for the degradation and breakdown of numerous extracellular matrix components such as collagen, leading to the degradation of cartilage (Burrage *et al*, 2006; Yoshihara & Yamada, 2007). The synergistic activity of these factors (RANKL and MMPs) leads to the gradual degradation of bone and cartilage in the joint area, leading to stiffness, pain and eventually disability of movement. Lastly, RA-FLS aberrant proliferation contributes to pannus formation and joint destruction. RA-FLS are the critical cell types in the growth of pathological synovial tissue in RA, and inhibition of their proliferation is a potential antirheumatic therapy (Sandler *et al*, 2006). RA-FLS are shown to be apoptosis-resistant (Zhao *et al*, 2021). The intrinsic and extrinsic pathways regulate apoptosis. The intrinsic pathway is activated inside the cell following intracellular stress or injury, including mitochondrial proteins like BH3 interacting domain death agonist (BID), Bcl-2-associated X protein (BAX), B-cell lymphoma 2 protein (BCL2), and phorbol-12-myristate-13-acetate-induced protein 1 (PMAIP1), among others. The tumour protein P53 (TP53) transcription factor is a critical activator of the intrinsic pathway of apoptosis via activating mitochondrial proteins. TNF and FAS ligand (FASLG) mainly activate the extrinsic pathway. Ultimately, both pathways activate caspases, which initiate a proteolytic cascade leading to cellular death (Peter, 2011).

The biological information regarding pathways implicated in the RA FLS pathology has been assembled into a large-scale mechanistic network, the RA map (Singh *et al*, 2020). While the RA map contains generic information about pathways implicated in RA that come from various sources, the RA FLS is the dominant cell type. The map is a process description diagram (Rougny *et al*, 2019) and is part of the Disease Maps initiative (Mazein *et al*, 2018; Ostaszewski *et al*, 2019). This type of construct contains valuable disease-specific information encoded in human and machine-readable formats. Besides serving as templates for omic data visualisation, disease maps can also work as scaffolds for building mathematical models and sources of causal interactions (Touré *et al*, 2021).

Dynamical modelling has been widely used to study and decipher complex biological processes that are otherwise hard to comprehend. Previous attempts to build computational models for RA have contributed a few kinetic models to study the role of proinflammatory and anti-inflammatory cytokines (Baker *et al*, 2013) and bone erosion (Macfarlane *et al*, 2019), and also the behaviour of various cells, including RA-FLS in cartilage destruction in RA joints (Moise & Friedman, 2019). A hybrid mathematical modelling framework that describes pannus production in a tiny proximal interphalangeal (PIP) joint was also proposed (Macfarlane *et al*, 2022). Nevertheless, the need for kinetic parameters limits its use for large-scale molecular interaction networks. Lastly, a large-scale hybrid model covering signalling, gene regulation, and metabolism in RA-FLS was published (Aghakhani *et al*, 2022). The model focuses on the metabolic reprogramming of fibroblasts under hypoxic conditions in the arthritic joint and suggests a reverse Warburg effect as the origin of the observed metabolic switch.

Discrete logic-based qualitative modelling has been increasingly used to model large-scale networks for which kinetic data is scarce (Hemedan *et al*, 2022; Hall & Niarakis, 2021). For example, in a Boolean model, each node can take only two values, 0 (FALSE) and 1 (TRUE). The next value of each variable is determined by a logical function (using the classical AND, OR, NOT operators) of the current values of its regulators (upstream nodes). The evolution of each node also depends on the updating scheme chosen. The synchronous scheme updates all variables in the model simultaneously; in the asynchronous scheme, the variables are updated individually in a non-synchronous manner (Niarakis & Helikar, 2021; Abou-Jaoudé *et al*, 2016).

In this work, we use the state-of-the-art RA map (Singh *et al*, 2020) and the tool CaSQ (Aghamiri *et al*, 2020) to build a large-scale, modular Boolean model of RA-FLS focused on inflammation, bone erosion, cartilage destruction, cell proliferation, and apoptosis. The model consists of five phenotype-specific sub-models (to be mentioned as *modules* later in the paper) that can be simulated individually and a five-phenotype global model (to be mentioned as the *global model* later in the paper). Systematic testing of different initial conditions using the modules and the global model showed that both the modules and the global model could reproduce small-scale experimental results from the literature. To search for meaningful steady states, we reduce calculations by propagating and eliminating fixed input components, as introduced by (Saadatpour *et al*, 2013) and implemented in the CoLoMoTo notebook (Naldi *et al*, 2018) by (Hernandez *et al*, 2020). Input propagation consists of assigning fixed values to some of the model’s inputs and using Boolean algebra to simplify the rules of their downstream components. We also use a probabilistic framework to calculate phenotypic probabilities starting from predefined initial conditions. The RA-FLS model was used to study the effects of mono and combined therapy for RA and suggest potential targets that could enhance the desired phenotypic outcome. Furthermore, drug repurposing analysis identified possible drug candidates that have as targets the previously identified nodes. A new round of simulations was then performed to evaluate their impact on the cellular phenotype and, subsequently, on the arthritic joint.

## Results

### Enhancing the cell specificity of the initial network

RA-FLS is the most frequent cell type in the RA map, covering a total of 45%, followed by synovial tissue with 36% (Singh *et al*, 2020). We used the updated fibroblast overlay list provided by (Zerrouk *et al*, 2022, 2020) to calculate the RA-FLS specificity of the global model. In this study, they used the RA fibroblast list provided by (Singh *et al*, 2020) and updated it with the RNA-seq single-cell dataset available in the GEO database (Mizoguchi *et al*, 2018) by performing Differential Expression Analysis (DEA) using BioTuring software and the Venice method. Using this list, out of 261 unique model components (proteins, genes, phenotypes), 194 are RA-FLS specific. Other cell types primarily include synovial tissue with 29 components, peripheral blood mononuclear cells (PBMCs) with 25 components, along with the presence of other sources like blood, synovial fluid, T-helper (th1), macrophages and chondrocytes (**Supplementary Table 1**).

### Five phenotype-specific modules and global RA-FLS model

The tool bioLQM (Naldi, 2018) was used to extract the phenotype-specific modules and the global model using the SBML file produced by CaSQ (Aghamiri *et al*, 2020) using RA map XML file as input (files available on GitLab). The global model can be seen in **Fig 2**. The size of the models in terms of nodes and reactions can be seen in **Fig 3A**. One straightforward observation is that the size of the five phenotypes global model is not the sum of the subparts. Indeed, as seen in the Venn diagram (**Fig 3B**), there is a core of 191 nodes shared by all modules, and only a few nodes are characteristic of each phenotype-specific module.

**Figure 1.**
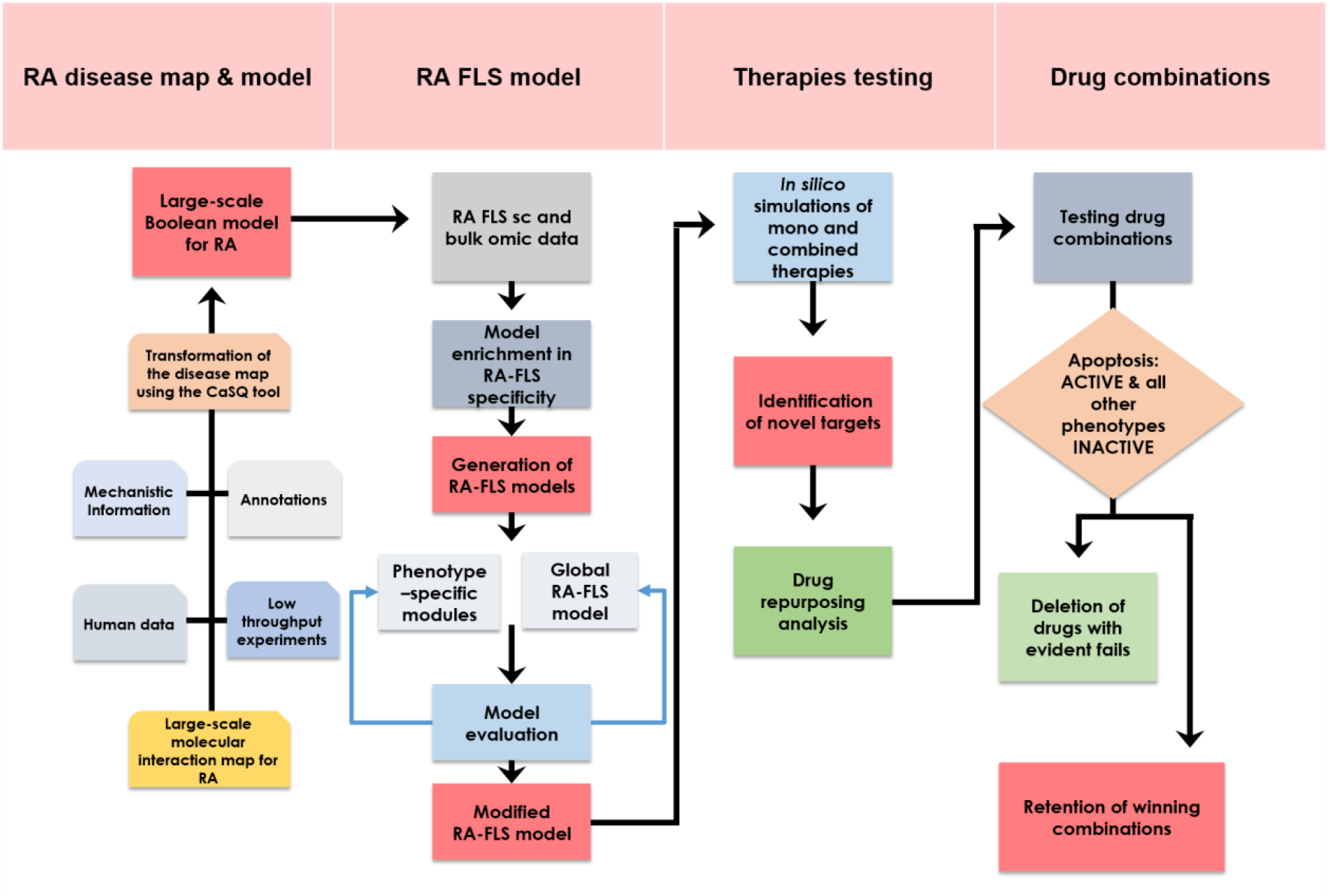
Construction and simulation of a large-scale, modular Boolean model of RA-FLS for evaluating novel drug combinations. The RA map was converted into an executable Boolean model using the map-to-model framework described in Aghamiri et al, 2020. Using single-cell omic datasets and literature studies, the RA generic model was subsequently enriched in RA-FLS-specific data. The RA-FLS model focuses on five phenotypes (apoptosis, cell proliferation, inflammation, matrix degradation, and bone erosion) characteristic of RA’s fibroblasts. Individual phenotype-specific sub-models and a five-phenotype global model were created. Biological scenarios extracted from the literature were used to evaluate and validate the models’ behaviour leading to some modifications of the original models. The modified RA-FLS model was then used to test mono and combined RA therapies. Drug repurposing analysis and further drug combination simulations led to a panel of suggestions of drug combinations that are predicted to have a favourable outcome (apoptosis active, cell proliferation, inflammation, bone erosion and matrix degradation inactive).

**Figure 2:**
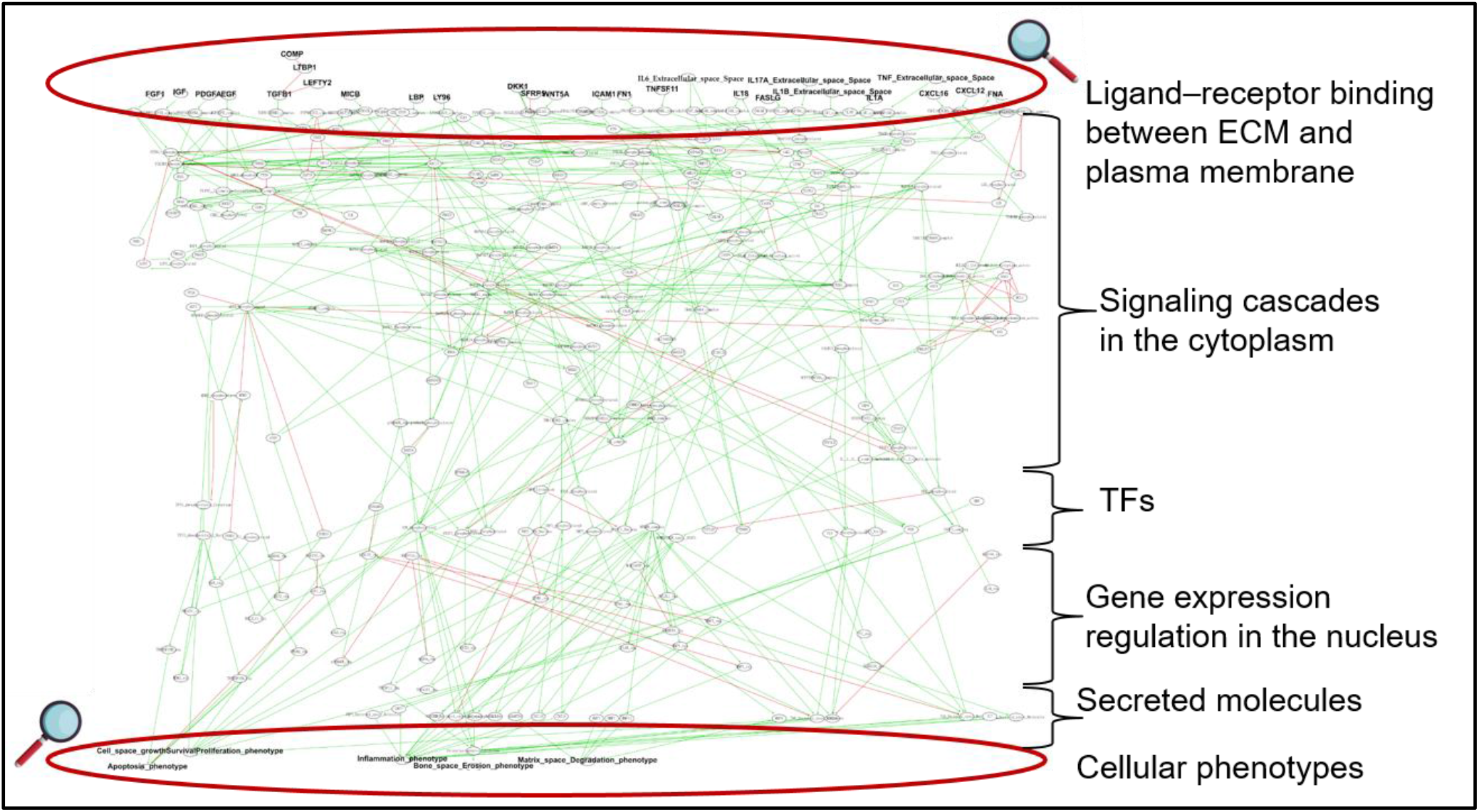
The regulatory graph of the RA-FLS global model. The layout is similar to the molecular map, with ligands/receptors and proximal signalling at the top, second messengers in the cytoplasm, gene regulation in the nucleus, secreted components and phenotypic outcomes at the bottom of the graph. Green arcs denote activation, and blunt red arcs denote inhibition. The regulatory graph comprised 321 nodes and 515 interactions (magnifying glass vector from vecteezy.com).

**Figure 3.**
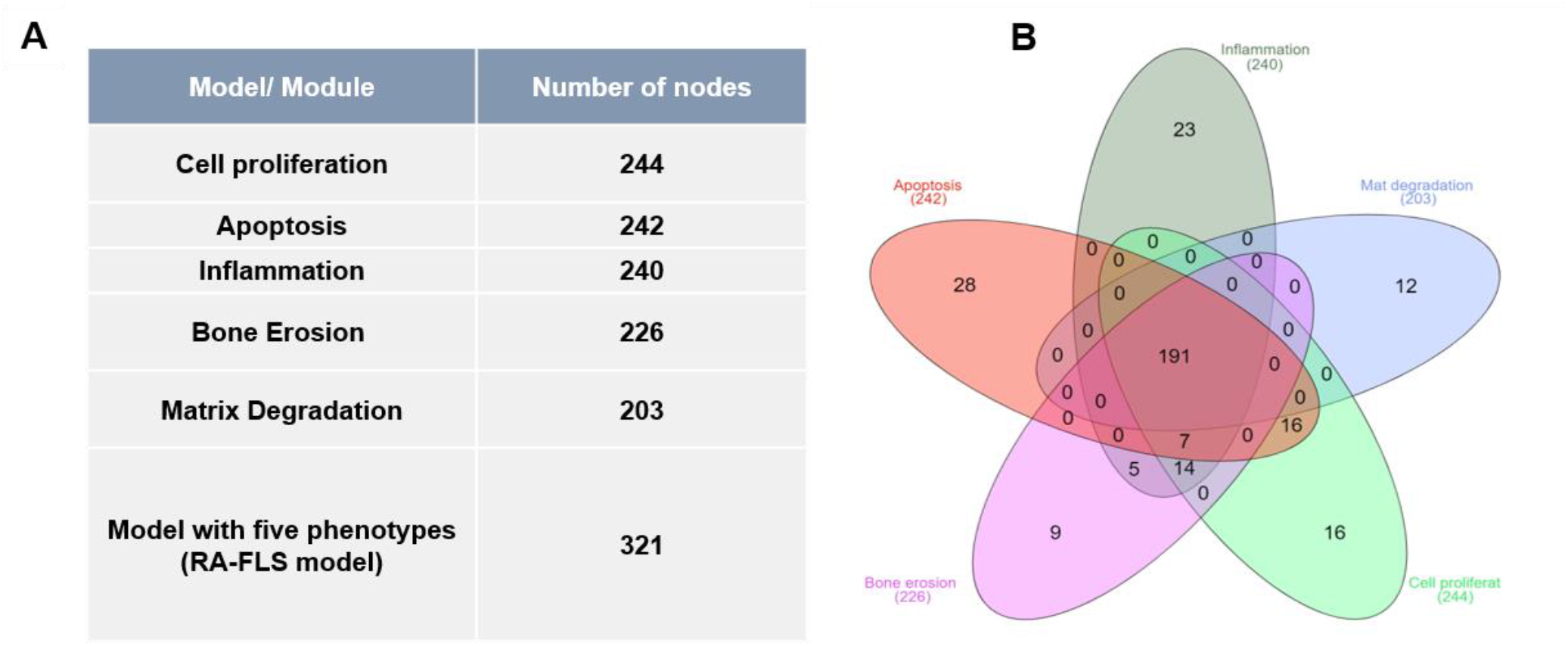
A) Number of nodes of the phenotype-specific modules and the global model. B) Venn diagram of all the five phenotype-specific components. The core of 191 nodes is shared among all five modules, and only a few are characteristic of the corresponding phenotype-specific module.

### Evaluating the RA-FLS model under different sets of input configurations using the input propagation method

The validation of each phenotype-specific module and the global model was performed by comparing the outcome of *in-silico* analyses to RA-FLS-specific biological observations based on experimental evidence. For each module, we formulated prior knowledge based on small-scale experiments as observations and compared them with the results of the corresponding virtual experiment. Initial conditions were set to mimic the corresponding experimental settings. We used input propagation and identification of trap spaces to evaluate the phenotype-specific modules and the global model’s behaviour. First, we assessed the model’s behaviour by comparing the intended/ expected biological behaviour and the observed one after the process. In general, we faced some difficulties in assessing the models’ behaviour that can be summarised in the following categories: a) not all components mentioned in the experimental settings included in the models, b) experimental information not available for all components included in the models, c) global behaviour consistent with experimental observations, but intermediate mechanisms only partially coherent. For some cases, further testing of the models revealed additional conditions needed to replicate the anticipated results. These additional “model conditions” that appeared to regulate the biological process are not experimentally proven (Supplementary Table 2).

#### Inflammation (Inflammation_phenotype in the models)

We formulated six biological scenarios to model Inflammation using the modules and the global RA-FLS model. TNF, one of the major pathways regulating Inflammation, was only able to activate the phenotype with the additional activation of the IKBA/NFKB/RELA complex. IKBA/NFKB/RELA complex is an input for activating the NFKB pathway, which is a key regulator of transcriptional responses to TNF (Matsuno *et al*, 2002; Hayden & Ghosh, 2014; Liu *et al*, 2017). IL-6 activation is a sufficient signal for activating Inflammation, as no additional inputs are needed to propagate the signal until the phenotype. Regarding IL-17, similar to TNF, the NFKB pathway is needed to activate the inflammation phenotype (Sønder *et al*, 2011; Hata *et al*, 2002; Liu *et al*, 2017).

#### Matrix degradation (Matrix_Degradation_phenotype in the models)

Cartilage destruction is one of the debilitating characteristics of RA. Activated RA-FLS have been shown to attach to the cartilage surface and release matrix-degrading enzymes. Matrix metalloproteinases (MMPs) play a pivotal role in cartilage destruction (van der Laan *et* al, 2003). Our model reproduced the destructive role of two MMPs, MMP1 and MMP9, in activating the matrix degradation phenotype in both the phenotype-specific module and the global model.

#### Bone erosion (Bone_Erosion_phenotype in the models)

Bone erosion is another significant characteristic of RA. Synovitis, along with the production of proinflammatory mediators like Wnt (wingless-related MMTV integration site) and receptor activator of nuclear factor κB ligand (RANKL), results in the differentiation of bone-resorbing osteoclasts, thereby stimulating local bone resorption(Cici *et al*, 2019). Secreted frizzled-related protein 5 (SFRP5), the primary upstream negative regulator of the WNT pathway when active, negatively regulates bone erosion (Chen *et al*, 2017). Trap spaces showed that with both the module and the RA-FLS model, we could reproduce Wnt and RANKL (in the presence of SFRP5) biological behaviour(Pettit *et al*, 2001; Kwon *et al*, 2014; MacLauchlan *et al*, 2017). Simulations of the models revealed a key role for SFRP5 in activating the bone erosion phenotype, as in its absence, the Wnt canonical pathway gets activated and further activates bone erosion.

#### Cell proliferation (Cell Growth/Survival/Proliferation_phenotype in the models)

Various growth factors like Platelet-derived growth factor (PDGFA) and transforming growth factor beta 1 (TGFB1), and cytokines like TNF regulate the proliferation of RA-FLS (Rosengren *et al*, 2010; Zhang *et al*, 2019). Platelet-derived growth factor (PDGF) is an essential mitogen for fibroblasts, including RA-FLS (Thornton *et al*, 1991; Remmers *et al*, 1991). PDGFA activated cell growth in both the phenotype-specific module and the global model. However, other pathways can still activate cell growth.

#### Apoptosis (Apoptosis_phenotype in the models)

The impaired apoptosis process of RA-FLS is responsible for synovial hyperplasia and joint destruction. Major pathways regulating apoptosis include extrinsic (FASLG and TNF) and intrinsic (mitochondrial pathway with BCL2 family proteins). Extrinsic pathways like FASLG and TNF contribute to the activation of apoptosis. The apoptosis module and the global model were able to reproduce this behaviour. AKT is another intracellular regulator of apoptosis (anti-apoptotic agent), which, when kept ON, protects RA-FLS against the apoptosis induced by FASL through inhibition of BID cleavage. However, this scenario was not reproducible, as discussed in the material and methods.

One of the reasons for the inconsistency was the presence of a direct inhibition (CAV1 negative regulator) to the apoptosis phenotype and the inferred logical formula that comprised OR gates between the activators and the absence of inhibitors. This direct inhibition reflects missing mechanistic details related to CAV1 regulating apoptosis. The RA map depicts this information using a negative influence directly at the phenotype glyph. While permitted by the CellDesigner tool (Funahashi *et al*, 2007), the lack of mechanistic details leads to an inferred rule from the tool CaSQ that is not optimal, as the tool is designed to use reactions (SBGN-PD) and not influences (SBGN-AF) as input. Therefore, to reproduce the apoptotic-resistant nature of RA FLS, we modified the logical formula changing the gates from OR to AND, keeping CAV1 as a dominant-negative regulator, in what we will denote the *modified global model*.

In our modified model, we need to have CAV1 set as active to act as a dominant inhibitor of apoptosis. This condition is justified and relevant from a biological point of view, as experiments have shown that MIR192 - which acts as a CAV1 inhibitor in our model, suppresses cell proliferation and induces apoptosis in human RA-FLS by downregulating caveolin 1. Moreover, MIR192 appears to be downregulated in RA synovial tissues and restoring its expression restores the growth-suppressive effects on RA-FLS by targeting CAV1 (Li *et al*, 2017). Therefore, the model condition of having CAV1 always active implies the absence of its inhibitor, MIR192, in the RA settings.

All analyses were performed for both versions of the RA-FLS global model, unmodified and modified, as seen in **Supplementary Table 2**, to ensure that the formula change did not have any impact on the model’s behaviour regarding the other four phenotypes. All biological scenarios in **Supplementary Table 2** were simplified and compiled into tables with the anticipated outcomes associated with Boolean values 0 (OFF) and 1 (ON). Value propagation analysis and trap space identification were applied, and heatmaps were used to visualise the results for both the global model and its modified version. We could not find experimental values for all the model inputs (88 inputs) in the RA condition. Therefore we opted to go for either 0 (no severe inflammation) or 1 (fully activated system, resembling more to an entirely inflammatory condition) for the remaining model inputs that were not part of the biological scenarios tested. In **Supplementary Figure 1**, we can see the behaviour of the global and modified model when all inputs are zero. All the phenotypes exhibit the same behaviour except for apoptosis. In the modified model, an additional condition of MIR192, a regulator of the negative inhibitor of apoptosis set as active, was required to reproduce two relevant scenarios.

We further analysed the modified model with input conditions as 1 (**Figure 4A**). Under these conditions, apoptosis remains OFF, and all other phenotypes remain ON, except for inflammation, which becomes inactive when IL6 is OFF. Furthermore, when all inputs resemble an inflamed state in these phenotypic behaviours, many biological scenarios were still reproduced (as seen in **Figure 4B**, represented with dark grey colour). Lastly, some conflicts were observed between the expected and the displayed behaviour, represented in yellow. These conflicts were resolved with additional biological or model conditions in the next row.

**Figure 4.**
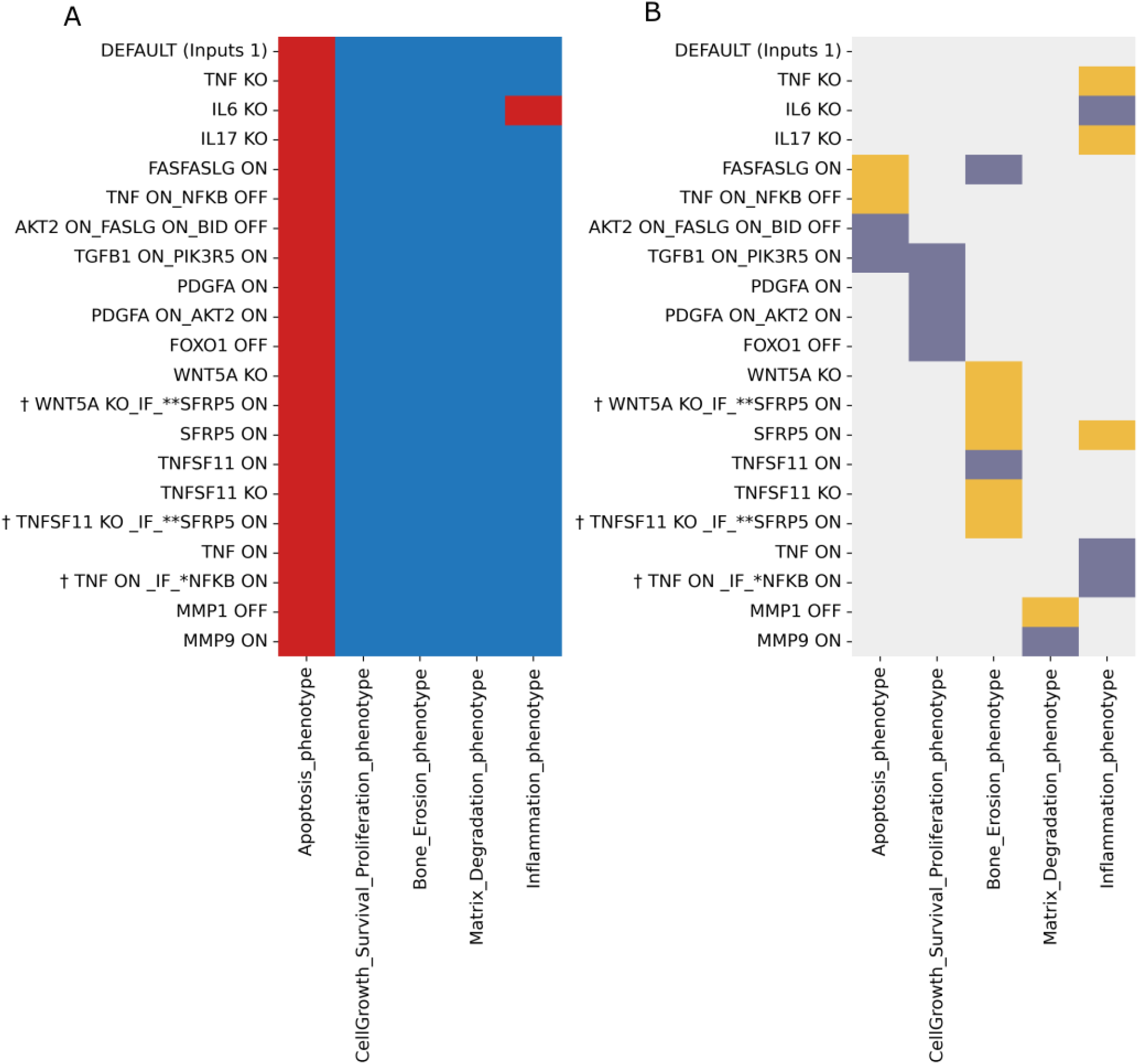
Heatmap displaying the trap spaces for the tested biological scenarios in the modified global model (A) and the comparison between expected and obtained values shown with colour codes (B). The y-axis shows all the tested scenarios’ names, as mentioned in table 2, regarding all the five phenotypes as outcomes on the X-axis with DEFAULT inputs set to one. ^**†**^ represent scenarios where additional conditions were given as a known biological behaviour* or as model conditions** **Trap spaces colour codes:** −1 (unfixed) 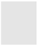, 0 (OFF) 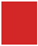, 1 (ON)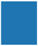 **Expectation graph colour codes:** **Score: expected value, obtained value; 1**: Yes [OFF, OFF & ON, ON] 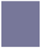; **0**: No [ON, OFF & OFF, ON (conflict)]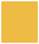, **-1**: Undefined 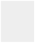

In general, evaluating the models’ performance in regards to more mechanistic processes and intermediates that involved more complicated scenarios was not possible in several cases, as experimental information is currently missing, and we could not conclude on the coherence of these results. This is reflected in **Figure 4B (and also Supplementary Figures 1B and 1D)** for the cases represented with light grey colour. Nevertheless, these scenarios could serve as interesting hypotheses for further experimental testing.

### Calculating continuous time phenotypic probabilities

The RA-FLS model, in both versions, was used to calculate phenotypic probabilities. We used the software MaBoSS to reproduce representative scenarios with different parameters (discussed in the Materials and Methods section) (Stoll *et al*, 2017). One scenario per phenotype from those listed in **Supplementary Table 2** was selected for simulations, and results showed that in all cases but apoptosis, the two model versions were able to reproduce the experimental observations. In brief, we performed the following simulations: ***Inflammation*** - IL6, one of the main pathways regulating inflammation, was set active (1), and the Inflammation phenotype was chosen as the output. As a result, the simulation confirmed the activation of inflammation (**Figure 5A**).

**Figure 5.**
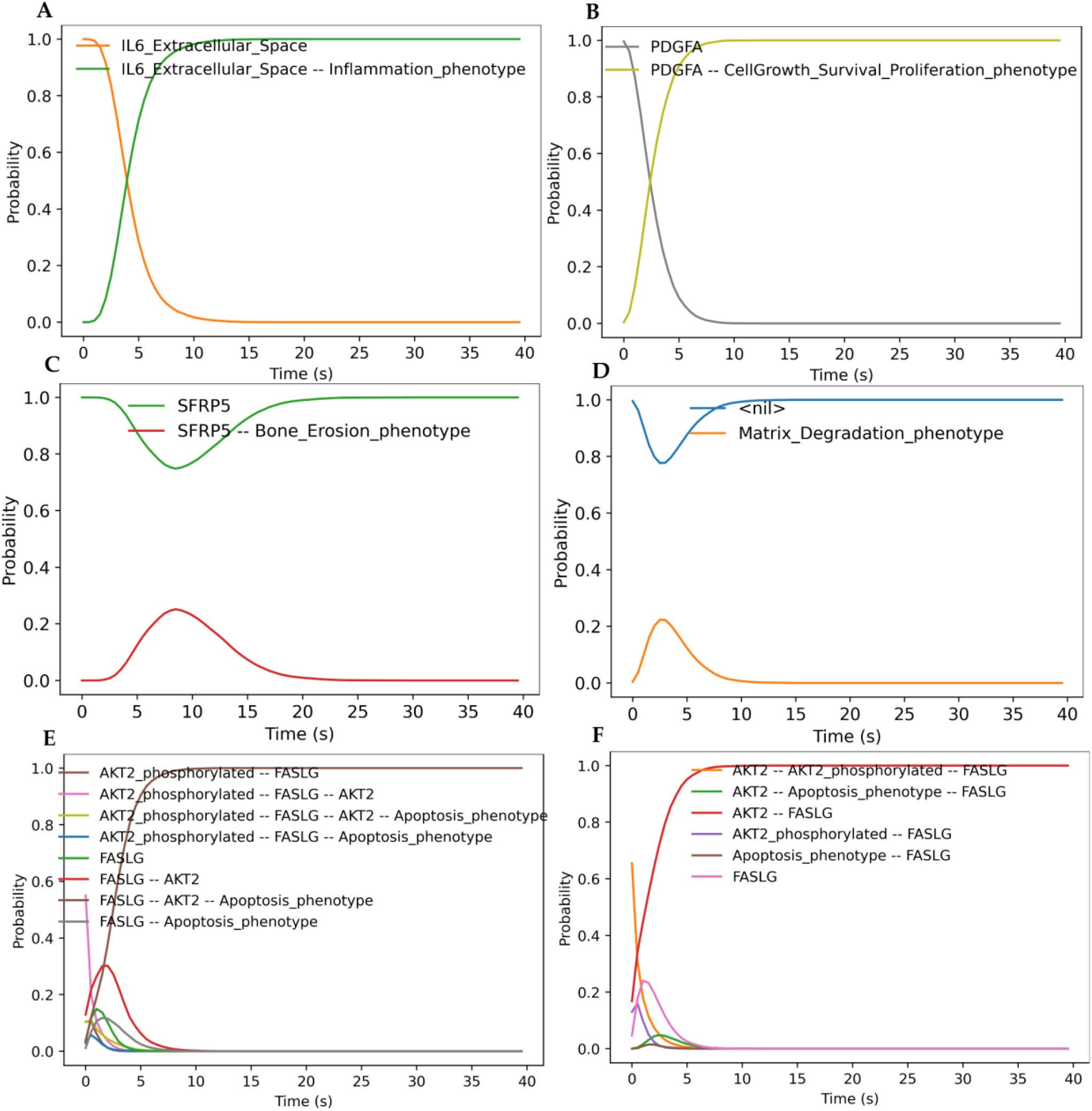
Calculating continuous time phenotypic probabilities of the selected initial conditions. **A)** Simulation with IL6_Extracellular_space active and Inflammation as output. The Inflammation phenotype gets activated in the presence of IL6 **B)** Simulation with PDGFA active and Cell proliferation as output. The Cell proliferation phenotype gets activated in the presence of PDGFA **C)** Simulation with TNFSF11 (RANKL) inactive, SFRP5 active and Bone erosion as output. The Bone erosion gets deactivated in the presence of SFRP5 and absence of TNFSF11 **D)** Simulation with MMP1 inactive and Matrix degradation as output. Matrix degradation phenotype gets deactivated in the absence of MMP1 **E)** Simulation with FASLG active, AKT2 active, BID inactive and Apoptosis as output (RA original global model). Apoptosis gets activated in the presence of FASLG, AKT2 and absence of BID **F)** Simulation with FASLG active, AKT2 active, BID inactive and Apoptosis as output (RA modified global model). Apoptosis gets inactive in the presence of FASLG, AKT2 and the absence of BID due to the dominant negative regulator CAV1.

#### Cell proliferation

When PDGFA was set as active (1), cell proliferation was activated (**Figure 5B**).

#### Bone erosion

When TNFSF11 (RANKL) was set as inactive (0) and the negative regulator SFRP5 as active (1), simulations showed the deactivation of bone erosion. (**Figure 5C**).

#### Matrix degradation

MMP1 was set as inactive by mutating its value as 0 (because it is an internal component and not an input to the model, its value must be fixed so that it cannot be changed by any upstream regulation, leaving it at 0 for the duration of the simulation) with matrix degradation chosen as the output. The simulation resulted in the deactivation of the matrix degradation (**Figure 5D**).

#### Apoptosis

FASGL and AKT2 were set as active (1) and BID inactive (0). In the unmodified RA-FLS global model, apoptosis was found to be active in this condition, while in the modified global model, apoptosis was found to be inactive due to the dominance of the CAV1 negative regulator (as seen in **Figures 5E, 5F**).

The results of the simulations are consistent with those from the trap space experiments for all scenarios tested.

### Finding trap spaces in extreme input conditions

The size and complexity of the modules and the global model make an exhaustive search for attractors in asynchronous mode prohibited. However, when starting from predefined initial conditions, attractors can be identified. For both versions of the RA-FLS global model, we searched for the trap spaces using two (extreme) input conditions: when all inputs are inactive (0) and when all inputs are active (1). We also chose some cytoplasmic (AKT2, JAK2, STAT1, CASP9, TP53, MDM2) and mitochondrial (BAX, BCL2) proteins as outputs, along with the five phenotypes. These proteins participate in the major intracellular pathways regulated by upstream external mediators and can serve as indicators of the activated pathways.

As seen in **Figure 6**, the only change observed between the original and the modified model concerns the apoptosis phenotype, which becomes OFF instead of unfixed when all inputs are OFF. This behaviour is biologically relevant as it matches biological evidence of fewer apoptotic cells in the joints of patients with active RA (Liu & Pope, 2003). It is worth noting that apoptosis remains ON in both models when all inputs are activated. This is because MIR192, an input of the model, is an upstream inhibitor of CAV1. CAV1 is, in turn, a dominant negative regulator of apoptosis; thus, when it gets inhibited by MIR192, apoptosis is ON. Apart from that, the behaviour of other model components, notably the mitochondrial pathway proteins BAX, BCL2, and CASP9, remained unchanged for both default conditions in both model versions. Regarding the cytoplasmic proteins, JAK2 and STAT1 remain inactive when the inputs are kept OFF and are activated once the inputs are ON, showing their direct dependencies on the regulation by the inputs and, more specifically, by IL6. Although the global behaviour of the model fits with the input-output expectation, reproducing the mechanistic details remains a challenge that requires a more in-depth understanding of pathway cross-talks.

**Figure 6.**
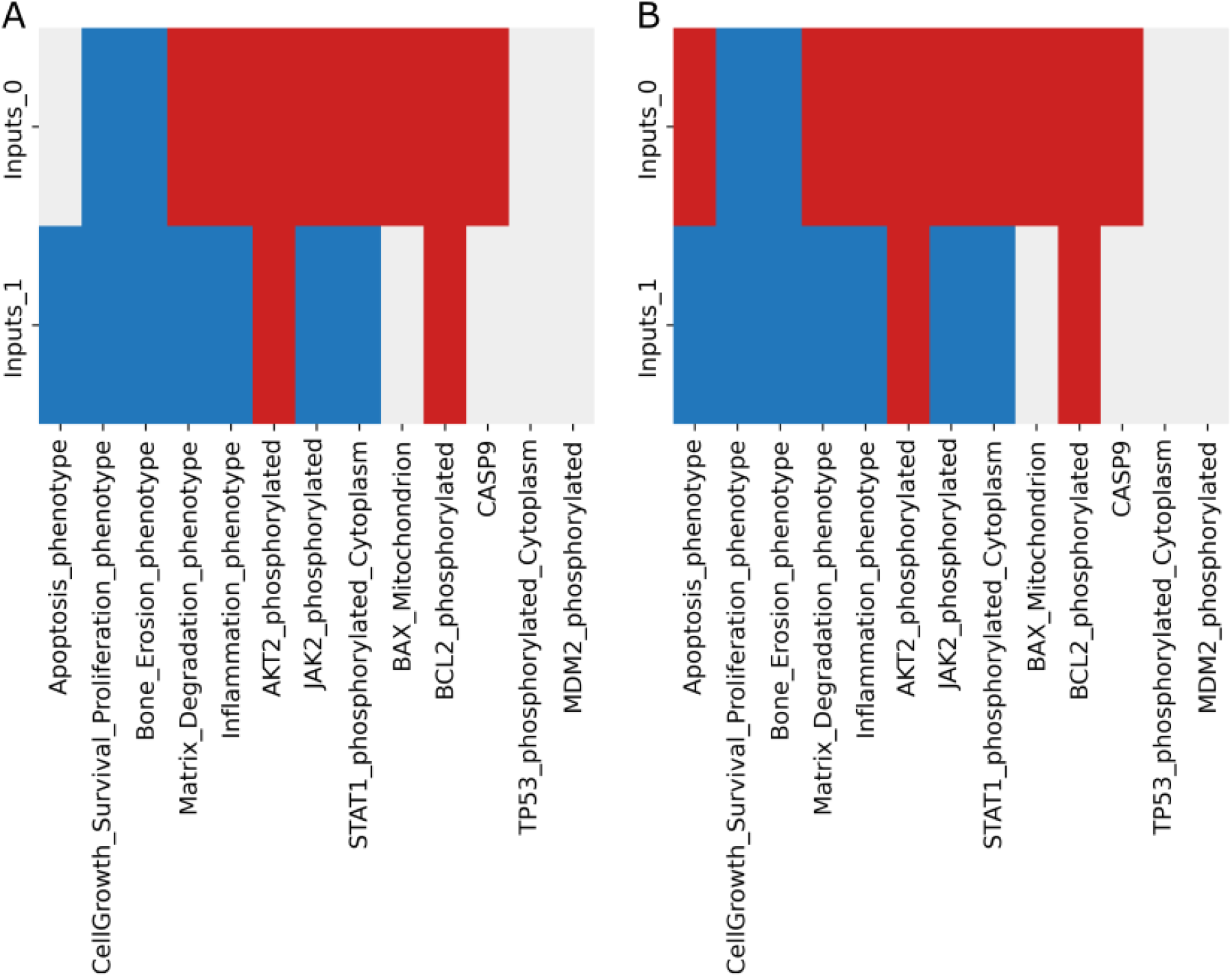
Trap spaces in extreme conditions with all inputs set as zero and one with the A) original global model and B) modified global model. Trap spaces colour codes: -1 (unfixed) 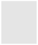, 0 (OFF) 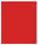, 1 (ON)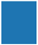

### Studying the role of p53 in the phenotypic outcomes of the RA-FLS model

The P53 protein is expressed in RA-FLS, and its overexpression is a characteristic feature of RA. This overexpression is probably induced by DNA strand breaks caused by the genotoxic environment of RA joints, in some cases because of P53 mutations. Independent studies from three groups indicated that p53 mutations occur in RA synovial tissue samples derived from a subset of RA patients. Furthermore, the inactivation of p53 may contribute to the invasiveness of FLSs and the high-level expression of cartilage degradation enzymes (Zhang *et al*, 2016; Sun & Cheung, 2002; Kullmann *et al*, 1999; Rème *et al*, 1998; Taubert *et* al, 2000). In summary, p53-dependent mechanisms could contribute to therapeutic outcomes depending on the type of intervention. As shown, the up-regulation of p53 may induce apoptosis and downregulate inflammation and proliferation.

On the other hand, the down-regulation of p53 may reveal key pathways involved in these mechanisms (Müller-Ladner & Nishioka, 2000). We used the modified version of the RA FLS model to perform relevant simulations. The model was imported to the web-based modelling platform Cell Collective (Helikar *et al*, 2012) and was simulated asynchronously. The simulation results for all inputs set as inactive showed that when TP53 was kept ON, apoptosis, matrix degradation and inflammation remained OFF, while bone erosion and cell proliferation remained ON. However, turning TP53 OFF under the same input settings had no impact on the unaffected phenotypic outcomes (**Supplementary Figure 2**). The same observation was made for simulations with all inputs set as active. Again, we observed that the turning ON or OFF of TP53 did not seem to influence the phenotypic outcome, as all phenotypes remained active regardless (**Supplementary Figure 3**). The results suggest that more complicated processes govern the RA FLS fate and that modifying the state of TP53 alone is not sufficient to change the cell fate decision.

P53 is also contributing to oscillatory behaviour through the P53 −MDM2 interactions. In short, a negative feedback loop on p53 is produced when p53 stimulates Mdm2 transcription, which in turn targets p53 for destruction. These P53 and MDM2 feedback-loop oscillations have been validated by mathematical and experimental models (Lev Bar-Or *et al*, 2000; Lahav, 2008) in various cell types and conditions. Moreover, TP53 directly takes part in the intrinsic apoptosis process by interacting with the multidomain members of the Bcl-2 family, causing mitochondrial outer membrane permeabilisation (Vaseva & Moll, 2009). We wanted to see if the modified RA-FLS model could reproduce this dynamic behaviour. First, we calculated trap spaces when all inputs were active. The trap spaces analysis showed that the values of the proteins BAX, CASP9, TP53, and MDM2 are unfixed, while those of proteins BAD and BID are fixed in an active state, and protein BCL2 is fixed in an inactive state (**Figure 7A**). The modified RA-FLS model was further analysed to study if the unfixed proteins exhibited an oscillatory behaviour under these conditions. We used the web-based modelling platform Cell Collective (Helikar *et al*, 2012) and asynchronously simulated the model setting all inputs as active. As seen in **Figure 7B**, the model can reproduce the oscillations between TP53 and MDM2 entities and in **Figure 7C**, we observe the oscillations in the intrinsic mitochondrial pathway of apoptosis.

**Figure 7.**
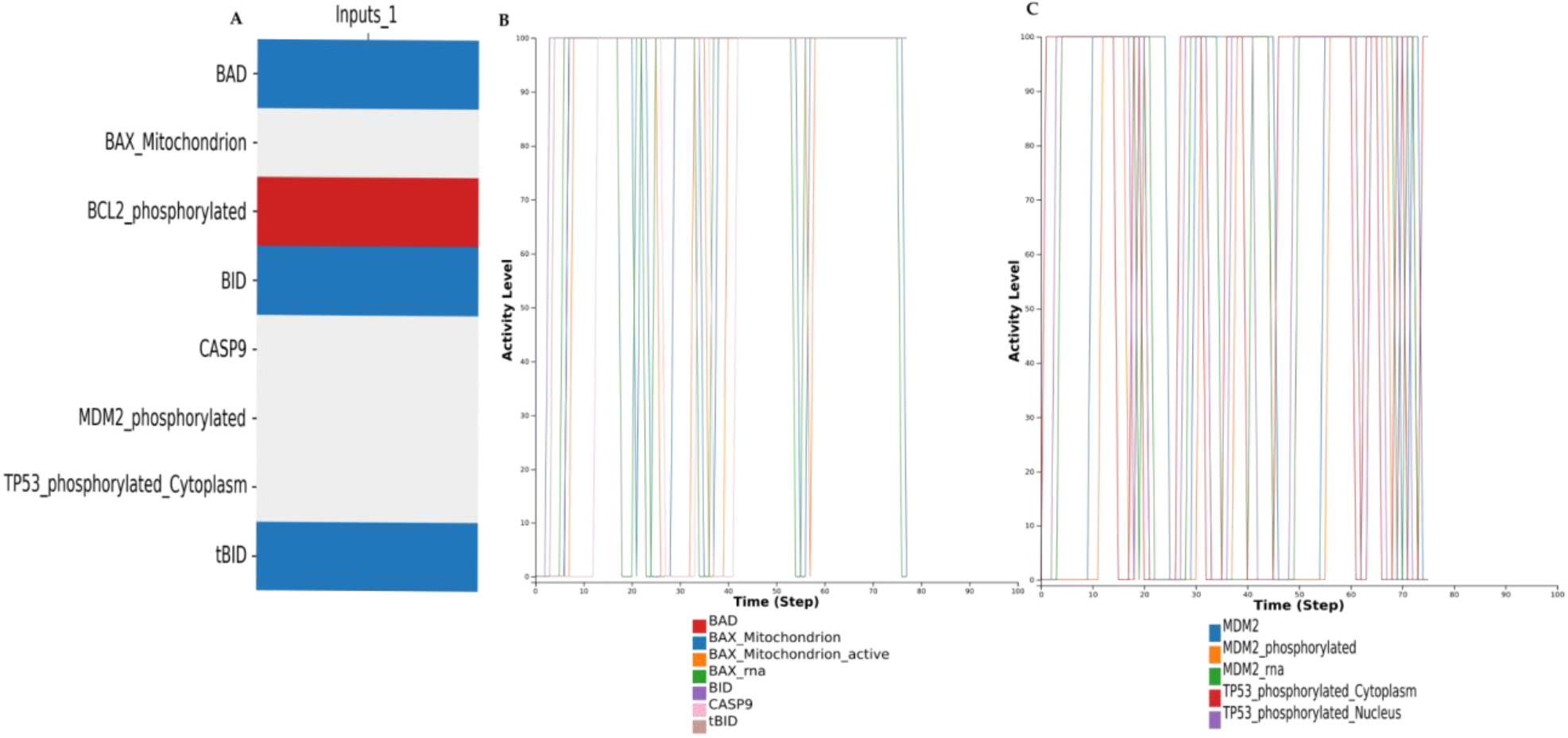
Oscillatory behaviour of TP53 and the mitochondrial proteins A) Trap spaces show that along with TP53 and MDM2, mitochondrial proteins like BAX and cytoplasmic protein CASP9 remain unfixed. B) Simulation with CC shows TP53 and MDM2 oscillations while all inputs remain active C) Simulation with CC shows oscillatory behaviour of the mitochondrial proteins when all inputs remain active.

### Reproducing mono and combined drug therapy in RA

Disease-modifying antirheumatic drugs (DMARDs), such as conventional synthetic (csDMARDs) (methotrexate, hydroxychloroquine), biologic (TNF-α inhibitors, IL-6 inhibitors), and targeted synthetic medications(tsDMARDs) (pan-JAK- and JAK1/2-inhibitors), have provided the most encouraging outcomes for the treatment of RA (Radu & Bungau, 2021; Lin *et al*, 2020). Many drugs targeting TNF and IL6 are already established in clinical treatment (Bullock *et al*, 2018). While the symptoms of inflammation and pain could be relieved with the cocktails of these medications, 30-40% of the patients fail to fully respond to treatment and experience periods of disease remission and relapse (Haschka *et al*, 2016). In addition, the side effects caused by most medications and the financial cost limit their use after a specific dosage (Wang *et al*, 2018; Lin *et al*, 2020). In **Table 1**, we provide a list of the mono and combined drug therapies used to treat RA. The RA-FLS model can mimic the effects of the drugs and predict the outcome of combined perturbations.

Analysis was performed with the modified version of the RA-FLS model, DEFAULT input conditions as 1, as they are closer to the inflamed conditions in the RA joint. In addition, we set CAV1 as 1 to reproduce the apoptosis-resistant phenotype of the cells. Results shown in **Figure 8** confirm that IL6, targeted by Sarilumab, Tocilizumab, and JAK, targeted by Tofacitinib, Baricitinib and Itacitinib, were able to downregulate inflammation. Under these conditions, all other phenotypes, except apoptosis, were found to be active and not impacted by the drug treatment, implying that their regulation is more complex and requires additional targeting. We also tested drug combinations that can be administered to RA patients. The combination of Methotrexate and Sarilumab was seen to deactivate inflammation successfully. Our results corroborate the experimental findings of clinical trials that demonstrated better efficacy of the sarilumab targeting IL6, either as monotherapy versus adalimumab, which targets TNF or in combination with csDMARDs versus placebo and csDMARDs (Genovese *et* al, 2020). Indeed, in our modelling framework, all results with anti-TNF drugs were inadequate to entirely suppress the inflammation phenotype, suggesting that blocking TNF alone is insufficient to hamper the signal. Our results also follow the findings of (Miagoux *et al*, 2021), which demonstrated that IL6 suppression was more successful in downregulating key transcription factors leading to inflammation in patients treated with anti-TNF treatment.

**Table 1.** Mono and combined drug therapy in RA.

| **Mono Drug therapy** | **Targets** |
| --- | --- |
| Tocilizumab, Sarilumab | IL6 (Huang *et al*, 2021) |
| Etanercept, Infliximab, Adalimumab, Golimumab, and Certolizumab Pegol | TNF (Huang *et al*, 2021; Bullock *et al*, 2018) |
| Tofacitinib (RA-FLS), Baricitinib, Itacitinib | JAK (Huang *et al*, 2021; Rosengren *et al*, 2012) |
| Secukinumab | IL17 (Huang *et al*, 2021) |
| Andecaliximab, Celastrol (RA-FLS) | MMP9 (Huang *et al*, 2021; Li *et al*, 2013; Fang *et al*, 2017) |
| Imatinib | PDGF (Sandler *et al*, 2006; Terabe *et al*, 2009) |
| Methotrexate | IL1B, PDGF, NFKB (Bergström *et al*, 2018; Spurlock *et al*, 2015) |
| Anakinra | IL1B (Mertens & Singh, 2009) |
| **Combined drug therapy** | **Targets** |
| Methotrexate + Sarilumab (or Tocilizumab) | NFKB, PDGFA, IL1B targeted by Methotrexate + IL6 targeted by Sarilumab (Boyapati *et al*, 2016; Genovese *et al*, 2015) |
| Methotrexate + Cyclosporine | NFKB, PDGFA, IL1B targeted by Methotrexate + Calcineurin targeted by Cyclosporine (Tugwell *et al*, 1995; Fox *et al*, 2003) |
| Methotrexate + Azathioprine | NFKB, PDGFA, IL1B targeted by Methotrexate + RAC1 by Azathioprine (Willkens & Stablein, 1996; Willkens *et al*, 1995) |
| Methotrexate + Hydroxychloroquine | NFKB, PDGFA, IL1B targeted by Methotrexate + TLRs targeted by hydroxychloroquine (Schapink *et al*, 2019; Carmichael *et al*, 2002) |
| Methotrexate + Infliximab or Golimumab | NFKB, PDGFA, IL1B targeted by Methotrexate + TNF targeted by Infliximab/Golimumab (Inui & Koike, 2016) |
| TNF inhibitor + Abatacept | TNF + CD80*, CD86*, CD28 targeted by Abatacept (Boleto *et al*, 2019; Korhonen & Moilanen, 2009) |
| TNF inhibitor + Anakinra | TNF + IL1 targeted by Anakinra (Boleto *et al*, 2019) |
| Abatacept + Anakinra | CD80*, CD86*, CD28 targeted by Abatacept + IL1 targeted by Anakinra (Boleto *et al*, 2019) |

\* absent in the model

**Figure 8:**
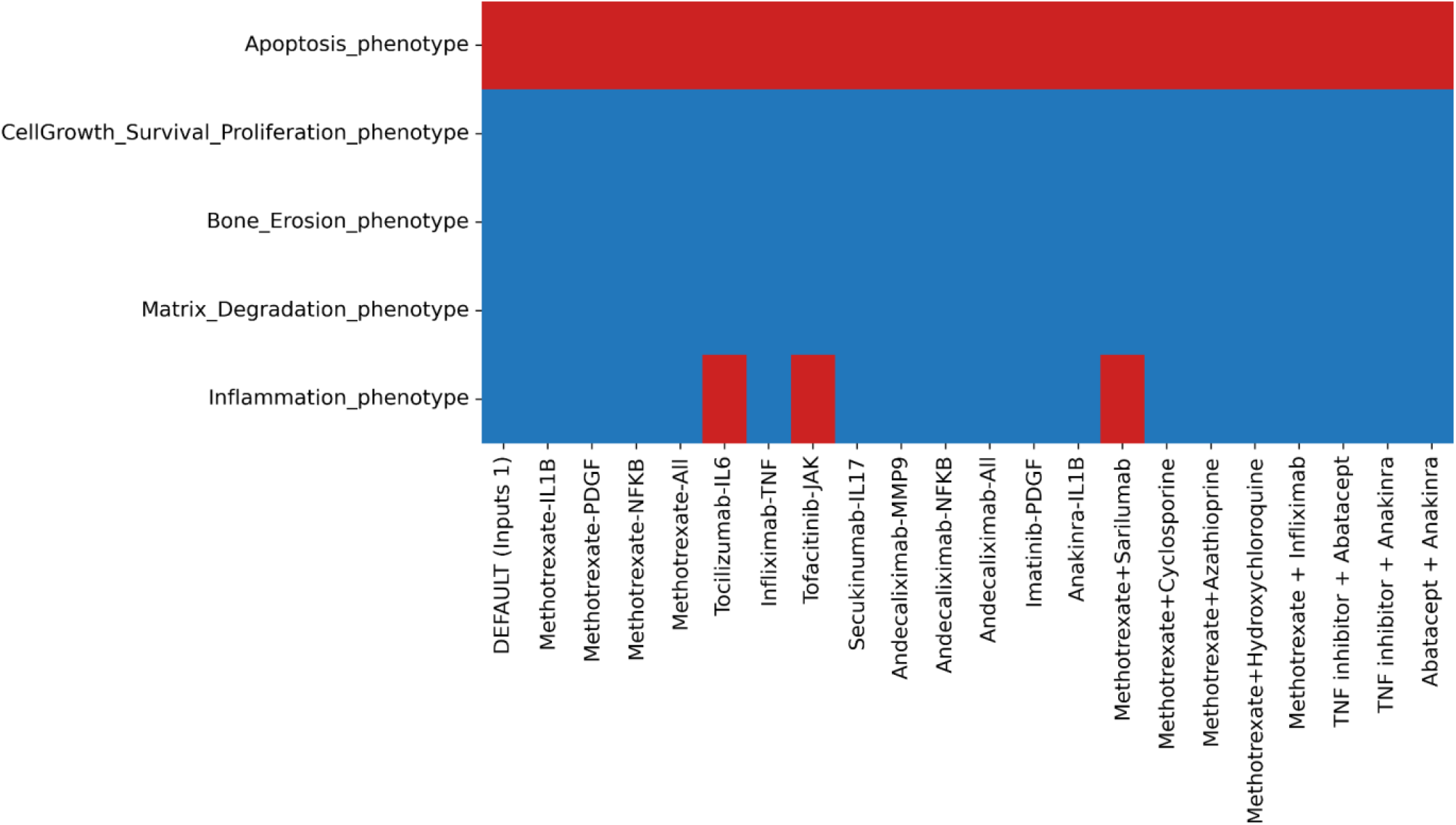
Heatmap showing identified trap spaces regarding the five phenotypes while testing different drug targets with DEFAULT inputs conditions as 1 with the modified version of the RA-FLS model.

### Identification of novel targets and candidate drugs via drug repurposing analysis

The main objective of our modelling study was to find conditions that can induce apoptosis and downregulate cell proliferation, inflammation, matrix degradation and bone erosion in RA-FLS. This included identifying targets and available drugs that could act upon those targets. To proceed, we decided to focus on the direct upstream regulators of each phenotype, study their role based on the logical formula that describes their effect on the phenotype, and search for the available drugs in clinical trials and *in vivo* and *in vitro* studies. The available drugs targeting the upstream regulators of each phenotype can be seen in **Supplementary Table 3**.

In our model, one of the core conditions to reproduce RA-FLS behaviour is setting CAV1 as always active (being a negative regulator of apoptosis), implying a downregulation of its inhibitor, MIR192, which is also an input. This condition is sufficient to reproduce the RA-FLS resistance to apoptosis along with the activation of cell proliferation, inflammation, bone erosion and matrix degradation (**Supplementary Figure 4**). We kept the same default model conditions (all inputs active and MIR192 inactive) for the drug targeting simulations using the modified RA-FLS model and the Cell Collective simulation platform (Helikar *et al*, 2012).

According to the logical formula, apoptosis is activated when CAV1 is inhibited and at least one activator is present. Interestingly, the negative regulator of apoptosis, CAV1, can be targeted by third-generation bisphosphonates (BPs), as demonstrated in various studies, although not always RA-relevant. Incadronate inhibits CAV1 expression in PC3 prostate cells (Iguchi *et al*, 2006). Furthermore, studies on human epithelial fibroblasts show the negative impact of BPs on the expression of genes essential for their growth and differentiation at medium-low therapeutic doses in the long term (Manzano-Moreno *et al*, 2019; Jung *et al*, 2018). As a substitute for patients who did not respond to conventional medication or who did not have easy access to biologics treatment, BPs have also been tested in RA patients. More specifically, scientists studied intravenous pamidronate administration for refractory rheumatoid arthritis. Results showed that pamidronate infusions were beneficial for most patients, but the alleviation of symptoms did not last for more than six months (Salesi *et al*, 2012). More recently, Zoledronic acid (ZA) was tested in combination with Methotrexate (MTX) in 66 RA patients for its efficacy in inhibiting RA disease activity. The combined treatment of ZA and MTX effectively reduced disease activity, fracture risk and bone pain in patients with RA-derived secondary osteoporosis (Xie *et al*, 2019). A clinical trial involving 28 patients to assess the effects of ZA on patients with early-stage RA and low disease activity was concluded in 2018, but the results are not yet available^1^. ZA has also been used to induce cell apoptosis in human and murine osteoclast precursors and mature osteoclast-like cells by triggering ROS- and GSK-3β-mediated Mcl-1 down-regulation (Tai *et al*, 2017). According to our model, BPs could also alleviate RA symptoms and inflammation, as they could inhibit CAV1 and activate apoptosis in RA-FLS. The inhibitory effects of ZA on tumour-related growth factor IL-6 in hormone-resistant prostate cancer cell lines have also been studied. The study showed the inhibitory effect of ZA on IL6 secretion, which in turn resulted in increased apoptosis (Asbagh *et al*, 2008). Keeping the default model conditions, we performed the simulations targeting CAV1 with Zoledronic acid. **The results showed that when CAV1 is inhibited, apoptosis will become activated** (**Supplementary Figure 5**).

Inflammation in RA is one of the most characteristic symptoms. The inflammation phenotype in the RA FLS model has thirteen direct upstream regulators. Among them, we find TNF, IL6, IL17A, IL1B, and NFKB, which represent common RA targets, and chemokines, such as CXCL8, CXCL9, CXCL10, and CXC11, two interferon proteins IFNB1, IFNA1, and lastly IRF1, IRF5 and IRF7. From al the direct upstream regulators, IRF1,5 and 7 seem to control the fate of phenotype and are all regulated upstream by the IL6 pathway. **We performed simulations under the default conditions targeting IL6, resulting in the complete inactivation of inflammation (Supplementary Figure 6**).

Bone erosion is directly dependent on the activation of eight upstream regulators. Besides TNF, IL17A, and IL1B, known targets of RA therapy, we also find IL7, TNFRSF11, FOS, JUN and NFATC1 to exert direct control over the phenotype. We were able to find drugs that inhibit all five of them, with one having been in clinical trials for patients with multiple sclerosis (MS), but based on misrepresentation of preclinical data (GSK2618960 for IL7). **Results showed that simultaneous blocking of IL7 and AP-1 (JUN and FOS) leads to the inactivation of bone erosion under the default model conditions (Supplementary Figure 7)**.

Matrix degradation in the RA FLS model seems to be controlled exclusively by zinc-dependent endopeptidases, namely four MMPs and a disintegrin and metalloproteinase with thrombospondin motif (ADAMTS) member, ADAMTS4. Among these proteins, MMP3 is the one that exerts complete control over the phenotype for the given default conditions, and **targeting only MMP3 was sufficient to inhibit matrix degradation** (**Supplementary Figure 8)**.

Regarding cell proliferation, eight direct upstream regulators can affect the phenotype, namely TNF, NFKB, P38, BCL2, RPS6KB1, ADAMTS9, CREB1, and YWHAQ. TNF and NFKB are both usual targets of RA therapy. Regarding NFKB, Bortezomib is a proteasome inhibitor that has been shown to downregulate its pathway in cancer cells (Sung *et al*, 2008). Bortezomib was also shown to attenuate murine collagen-induced arthritis (Lee *et al*, 2009) and was recently found to improve the joint manifestations of rheumatoid arthritis in three patients (Lassoued *et al*, 2019). P38 was once considered a promising target for developing new anti-inflammatory drugs to treat RA and other inflammatory diseases. However, the results in clinical trials were disappointing (Canovas & Nebreda, 2021). We identified drugs for two more upstream regulators, namely BCL2 and CREB1, and drugs that could target FOXO1, the upstream transcription factor of YWHAQ. **In default conditions, among the eight regulators, CREB1 and YWHAQ were shown to inhibit completely the cell proliferation phenotype when targeted (Supplementary Figure 9)**.

To recapitulate, the RA-FLS model was used to predict the best combinations for activating apoptosis and inhibiting inflammation, cell proliferation, bone erosion and matrix degradation. According to the model, the desired effect can be achieved by blocking CAV1, IL6, IL7 and AP-1 (JUN and FOS), CREB1 and YWHAQ, and MMP3. For blocking CAV1, we have three potential drugs, Pamidronate, Incadronate and Zoledronic Acid (ZA), with the latter having been tested for RA (**Supplementary Table 3**). Regarding IL6, Tocilizumab and Sarilumab are common biologics administered to RA patients (**Table 1**). IL7 can be targeted with Certolizumab pegol, GSK2618960, and CYT107, with Certolizumab already tested in RA patients (**Supplementary Table 3**). Concerning AP-1, we identified Acitretin and T-5224. Acitretin has been tested in clinical trials for various diseases. In psoriasis, it was also used in a small cohort of patients in combination with Etanercept (anti-TNF) to improve response to therapy, with encouraging results (**Supplementary Table 3**). For CREB1, we could only identify one inhibitor, 666-15, while for YWHAQ, we identified AS1842856, which can target and inhibit FOXO1, an upstream regulator of YWHAQ. However, both inhibitors have only been tested in mice studies. Lastly, for MMP3, a variety of broad-spectrum MMP inhibitors are available. Some have already been tested in RA patients (like Cipemastat or Apratastat) but failed to produce significant results. Most drugs targeting MMPs, in general, and MMP3, in particular, have been tested primarily on cancer disease **(Supplementary Table 3)**.

## Discussion

In this work, we focused mainly on the mechanisms regulating inflammation, matrix degradation, bone erosion, apoptosis, and cell proliferation, in RA FLS. Our framework presents the experimental and clinical data and model results in an approachable manner which links various software applications and data. To build our large-scale model, we used the state-of-the-art RA map (Singh *et al*, 2020) and the map-to-model framework proposed by Aghamiri et al, 2020 (Aghamiri *et al*, 2020). Five phenotype-specific modules and a global model were extracted and further enriched using RA-FLS-specific literature and single-cell data. The inferred models were further analysed using a variety of software, such as GINsim (Chaouiya *et al*, 2012), bioLQM (Naldi, 2018), MaBoSS (Stoll *et al*, 2012), and the CoLoMoTo notebook (Naldi *et al*, 2018). To our knowledge, the RA-FLS model is the first large-scale model that describes these cells’ signalling and gene regulation mechanisms in disease-specific settings. To evaluate the model’s behaviour, we mined experimental evidence based on cell-specific small-scale studies (knockouts and knock-ins) from the literature. We used value propagation (Saadatpour *et al*, 2013; Hernandez *et al*, 2020), trap spaces computation (Zañudo & Albert, 2013; Klarner *et al*, 2014), asynchronous simulations (Helikar *et al*, 2012, 2013), and continuous time Boolean stochastic modelling (Stoll *et al*, 2012) to analyse the models’ behaviour and evaluate the results. The employed methodologies not only reduced the amount of time needed to evaluate the behaviour of our large-scale models but also provided the means to comprehend intricate outcomes under specific circumstances.

The RA FLS model has successfully reproduced various experimental observations impacting the five phenotypes of interest. However, inconsistencies were also identified. For example, while working on apoptosis, we observed the inability of the apoptosis phenotype to be inactive in the presence of a negative regulator. It highlighted the insufficient knowledge related to the biological processes involved in regulating the phenotype and the limitations of the automated model inference. To leverage the missing information, we modified the logical formula making the inhibitor dominant, a choice also justified by experimental evidence. The modified RA FLS model reproduces many biological scenarios based on experimental evidence, at least in the input-output scale and for most of the phenotypes tested.

Furthermore, it successfully reproduces known oscillatory behaviour regarding TP53 and MDM2 along with the mitochondrial pathway proteins. The modules and the global model helped to understand the input and output (phenotype) relationships in a rather extensive network, along with the understanding of the regulatory processes at different biological scales. Considering it as a first step for understanding the RA-FLS behaviour in different sets of initial conditions, the model contributes toward a better understanding of the mechanisms that drive cell proliferation, inflammation, and resistance to apoptosis, bone erosion, and matrix degradation.

We used the RA-FLS model to perform mono and combined drug therapy simulations using drugs and combinations already administered to RA patients to understand better each drug’s different mechanisms of action and the combined effects on the cellular phenotypic outcomes. With all inputs active and mimicking the inflamed joint, IL6 produces a stronger signal for inflammation, as TNF and IL17 require the additional activation of NFkB to regulate inflammation. Additionally, while different DMARDs have distinct modes of action, they interfere primarily with the main pathways of the inflammation cascade (Benjamin *et al*, 2022). The activation of cell proliferation during drug testing and drug combinations shows that additional growth factors and signalling pathways are involved in controlling this process. As PDGFA is the only growth factor targeted in this instance, it seems insufficient to exert substantial control over the phenotype. Regarding bone erosion, the WNT non-canonical pathway signalling protein RAC12 (targeted by Methotrexate and Azathioprine) exerts partial control over the phenotype. However, bone erosion can also be triggered by other routes like RANKL (TNFSF11) and the WNT canonical pathway, thus keeping bone erosion active.

The next step was to identify possible targets that could enhance the desired effect of the treatment. Our goal was to achieve activation of apoptosis, suppression of inflammation, bone erosion, matrix degradation and cell proliferation for RA-FLS. To do so, we identified the direct regulators of the five different phenotypes and used drug repurposing analysis to identify drug candidates for these targets. More specifically, we used the logical formulae of the Boolean model to identify the direct regulators of each phenotype that could change the phenotypic state if targeted. We used several databases and dedicated sites to infer information about drugs and small molecules that target the action of these regulators. Some of these drugs had already been tested in RA or were administered in the context of other diseases. We had to tag discontinued drugs due to side effects, even when administered in a different disease context. For some drugs, information on toxicity or trial reports was limited. For example, CTS-1027, an MMP inhibitor, had been in phase II clinical trials by Conatus Pharmaceuticals (licensed from Roche) to treat Hepatitis C Virus (HCV) infection. However, this study was discontinued due to abnormalities and adverse events in a subset of clinical trial participants. This compound, originally discovered by Roche, showed positive results for osteoarthritis treatment, but no development report is available for this study^2^. Similarly, Marimastat, a broad MMP inhibitor, was discontinued in phase II trials, as it demonstrated drug toxicity linked to musculoskeletal pain and stiffness initially involving small peripheral joints of the hands (Keystone, 2002; Rasmussen & McCann, 1997).

While our framework can be used to identify potential drugs and drug targets, we cannot assess toxicity or drug dosage but suggest new potentially interesting drug combinations to be tested experimentally. In Table 2, we recapitulate the identified drugs for the highlighted targets. Combining them could help activate apoptosis and downregulate inflammation, matrix degradation, bone erosion and cell proliferation in RA-FLS.

**Table 2:** List of drugs found to upregulate apoptosis and downregulate inflammation, bone erosion, matrix degradation, and cell proliferation, targeting the identified biomarkers revealed by the RA FLS model.

| Identified drugs | Target components | Target phenotypes and expected effect |
| --- | --- | --- |
| Pamidronate, Incadronate, and Zoledronic Acid | CAV1 | Apoptosis ↑ |
| Sarilumab, Tocilizumab | IL6 | Inflammation ↓ |
| Certolizumab pegol, GSK2618960 and CYT107 and T-5224, Acitretin | IL7<br>AP-1 | Bone erosion ↓ |
| Batimastat | MMP3 | Matrix degradation ↓ |
| 666-15 and AS1842856 | CREB1<br>YWHAQ (FOXO1) | Cell proliferation ↓ |

## Limitations and perspectives

Our analyses demonstrated that besides input-output relationships, it is also necessary to understand and elucidate the underlying mechanisms that control the different phenotypes. Unit testing, wherein collections of tests specify the anticipated behaviour connected with distinct system modules, could be applied to verify local behaviour ((Saadatpour *et al*, 2013; Hernandez *et al*, 2020)). This could be of particular interest for understanding the complexity of the apoptosis mechanism in relation to the extrinsic and intrinsic (mitochondrial) pathways. Moreover, the resulting inconsistencies between prior knowledge and model behaviour could serve as a helpful indicator for re-evaluating the inferred logical formulae. The availability of more quality datasets could help to analyse the models under different sets of initial parameters and study their relation to the phenotypic outcomes. Furthermore, topological analysis of the model’s structural characteristics could reveal hubs and nodes with high centrality measures (Durón *et al*, 2019), which could guide the combination of *in silico* knockouts (KOs) and knock-ins (KIs) for simulating the models.

Our approach supports using systems modelling in preclinical and clinical settings to provide insights into the possible outcomes of drug combinations. Further optimisation of the RA-FLS model using patient-specific omic data integrated with other clinical parameters could help towards personalised treatments.

## Materials and methods

### Generation of RA-FLS phenotype-specific modules and the global model

Analysing complex biological networks often requires a workflow that includes several different tools with different specifications and requires many additional steps to harmonise the whole procedure. The CoLoMoTo Interactive Notebook provides an integrated environment to execute, share, and reproduce analyses of qualitative models of biological networks (Naldi *et al*, 2018). The framework is available as a Docker image on Linux, MacOS, and Microsoft Windows with the tools already installed and ready to be run with a Jupyter web interface executing the codes and visualising the results. The resulting notebook files can be shared and re-executed in the same environment. After installing the docker and accessing the latest colomoto-docker −V 2022-10-01 image, the notebook interface with python modules for different tools was available to execute the analysis.

A global, comprehensive and fully annotated RA-specific map was published recently (Singh *et al*, 2018, 2020). This map features components and interactions implicated in RA coming from various cell types. The RA-map CellDesigner XML file was first imported into the notebook using BioLQM and then converted into an executable RA-map model using CaSQ (CellDesigner as SBML-Qual), a tool for automated inference of large-scale, parameter-free Boolean models from molecular interaction maps based on network topology and semantics (Aghamiri *et al*, 2020). The executable RA model, in standard SBML-qual format (Chaouiya *et* al, 2013), can be further simulated and analysed using different modelling tools supporting the SBML-qual format. First, the RA model was sanitised using bioLQM to replace the model node Ids with node names. Then, the model was further used to extract five phenotype-specific modules, apoptosis, cell proliferation, inflammation, matrix degradation, bone erosion, and a global model. The modules were extracted by selecting the phenotype and extracting its upstream part of the model through the connected components. Within this framework, we developed python notebooks to validate RA-FLS-specific biological scenarios, drug testing and drug combination predictions.

### Model annotation and use of standards

Besides building and analysing large-scale biological models, one of the main challenges in systems modelling is the use of standards and proper annotation of such models to enhance a model’s reusability (Niarakis *et al*, 2022, 2021). The models obtained in our pipeline are in a standard format (SBML-Qual) and thoroughly annotated using the MIRIAM scheme (Le Novère *et al*, 2005), including PubMed IDs and pathway identifiers to annotate components and reactions. However, these elements are maintained and visualised only when the models are imported into the Cell Collective platform (Helikar *et al*, 2012) and lost when the models are further analysed with our CoLoMoTo notebooks due to the sanitisation process.

### Literature survey to find RA-FLS-specific biological scenarios

As our modelling efforts are based on RA-FLS, we did exhaustive literature mining to find biological scenarios to be tested for each phenotype with the individual modules and the global model. As a result, we formulated biological scenarios to be tested for each phenotype from the literature - 5 for inflammation, 5 for bone erosion, 4 for cell proliferation, 2 for matrix degradation, and 4 for apoptosis phenotype **(for more details, see Supplementary Table 2)**.

### Identification of trap spaces

We used *terminal trap spaces* to evaluate the asymptotic behaviour of our RA-FLS model. A *trap space* (also known as a stable motif or symbolic steady state) is a subspace from which the system cannot escape. (Zañudo & Albert, 2013; Klarner *et al*, 2014). In particular, all possible successors of every state of a trap space belong to the same subspace. A trap space is *terminal* if it does not contain any smaller trap space. Terminal trap spaces accurately represent the Boolean model’s asymptotic behaviour and provide a good approximation of all attractors in practice.

Trap spaces can be efficiently identified using constraint-solving techniques without performing the simulation. Here we used the implementation provided by the bioLQM software for the identification of terminal trap spaces in reduced models obtained after input propagation, as explained below.

### Propagation of fixed components

Value propagation works on the propagation of assigned constant values to the corresponding downstream nodes. First, the cellular context is defined by assigning constant values to some model components, followed by a model reduction technique (Saadatpour *et al*, 2013). Then, for each fixed (constant) node, the corresponding value is inserted into the logical rule associated with each target node. If the rule can then be simplified to a constant, this new fixed value is further propagated into the logical rules of downstream nodes. This process is iterated until no further simplification can be made on the logical rules of the model (Hernandez *et al*, 2020). Value propagation decreases the complexity of the model while preserving all attractors and terminal trap spaces.

### Continuous time stochastic simulations

MaBoSS is a C++ software for simulating continuous/discrete-time Markov processes applied to Boolean networks (Stoll *et al*, 2012, 2017). MaBoSS uses a specific language for associating transition rates with each node. Given some initial conditions, MaBoSS applies Monte-Carlo kinetic algorithm to the network to produce time trajectories and thus can associate probabilities with asymptotic solutions. The MaBoSS simulations were used to recreate at least one biological situation from each phenotype in the form of time-dependent (max time 40) trajectories using chosen parameters, more details of which could be found in the provided python notebooks.

### Simulations using the web-based platform Cell Collective

We also used the modelling platform Cell Collective to simulate the global models (Helikar *et al*, 2012). Models in Cell Collective can be imported via the SBML-qual standard (CaSQ-produced SBML-qual model file was imported) or built from scratch. The Cell Collective SBML-qual import is compatible with model annotations and network layout. Furthermore, references kept in CellDesigner’s XML file’s MIRIAM section can be retrieved and seen in the Cell Collective platform.

### Modified RA model

While validating the biological scenarios in the original model, we observed that while Apoptosis should have been OFF, the phenotypic node was always active. The inability of the phenotype to turn OFF was further examined by studying the regulatory rules, which indicated a lack of mechanistic information on the biological process. CAV1 was part of the OR rule for the apoptosis regulators, as can be seen below:

**!CAV1_rna**|**CASP3_phosphorylated**|**CASP8**|**TNFRSF10A_rna**|**TNFRSF10B_rna** where **‘**|**‘** represents a disjunction and **‘!‘** is a negation.

The Boolean rule was modified appropriately to add CAV1 as a dominant regulator:

**!CAV1_rna&(CASP3_phosphorylated**|**CASP8**|**TNFRSF10A_rna**|**TNFRSF10B_rna**, where **‘&‘** represents a conjunction.

This rule change generated a modified global model where apoptosis remains OFF in the presence of a dominant negative regulator corresponding to an actual RA condition where apoptosis remains deficient.

Five phenotype-specific individual modules (apoptosis, cell proliferation, inflammation, matrix degradation and bone erosion) and the global model were then extracted from the modified RA model. Finally, all the biological scenarios were again tested with the modified individual modules and the five phenotypes global model and compared with the unmodified modules and model results.

### Drug targeting (single, multiple targets) and drug combination analysis

We identified RA treatments that are administered regularly to RA patients and target different molecules, some of which are also RA-FLS specific (Table 2). We used both global models to simulate and analyse drug effects on the five selected phenotypes - Apoptosis, Cell proliferation, Inflammation, Matrix degradation and Bone erosion. We analysed the effect on the phenotypes for each drug by targeting its single and multiple targets in the global models. We also analysed the models by combining multiple drugs and their targets.

### Drug repurposing

We checked the logical formulae of all five phenotypes and extracted their upstream first-level regulators, as shown in **Supplementary Table 3**. For each regulator, we searched drugs with all clinical phases - launched, preclinical, phase 1, phase 2, and phase 3. For the latter, besides bibliographic search, the following sites were also used: https://clue.io/repurposing-app (Corsello *et al*, 2017); http://db.idrblab.net/ttd/ (Zhou *et al*, 2022); https://go.drugbank.com/ (Wishart *et al*, 2018). The results of this effort can be seen in **Table 2**, where the asterisk * denotes drugs already used in RA. We also thoroughly studied the mechanism of action of all the drugs (wherever information was available). We checked the default behaviour of the RA-FLS modified model concerning the five phenotypes by keeping all the inputs active and MIR192 inactive. MIR192 is input and needs to be OFF to activate the negative regulator of apoptosis CAV1, thus contributing to the state of RA pathology.

## Abbreviations

BID: BH3 Interacting Domain Death Agonist
BAX: Bcl-2-associated X protein
BCL2: B-cell lymphoma 2 protein
BPs: Bisphosphonates
CASP9: Caspase-9
FASLG: FAS Ligand
JAK1: Janus Kinase
1 IL-6: Interleukin 6
IL-1: Interleukin 1
IL-1B: Interleukin 1 Beta
IL-17: Interleukin 17
MDM2: Mouse double minute 2 homolog
MMPs: Matrix metalloproteinases
NFKB: Nuclear Factor Kappa-light-chain-enhancer of activated B cells
PBMC: Peripheral blood mono-nuclear cell
PDGFA: Platelet-Derived Growth Factors
PMAIP1: Phorbol-12-Myristate-13-Acetate-Induced Protein 1
RAC1: Ras-related C3 botulinum toxin substrate 1
RANKL: Receptor activator of nuclear factor kappa-B ligand
RA: Rheumatoid Arthritis
RA-FLS: Rheumatoid Arthritis Fibroblast-Like Synoviocytes
SFRP5: Secreted Frizzled-Related Protein 5
STAT1: Signal Transducer and Activator of Transcription 1
TGFB1: Transforming Growth Factor Beta 1
TH1: T helper 1 cells
TLRs: Toll-Like Receptors
TNF-α: Tumor necrosis Factor alpha
TP53: Tumor protein P53
Wnt: Wingless-related MMTV integration site
ZA: Zoledronic acid

## Data availability

All data is available in our GitLab repository - https://gitlab.com/genhotel/rheumatoid-arthritis-large-scale-computational-modeling/-/tree/main

## Conflict of interest

The authors declare no conflict of interest

## Authors’ contributions

ANi designed the study, supervised, performed simulation experiments, and analysed results. VS did literature search, contributed to notebooks’ development, calculated trap spaces, performed simulation experiments, analysed results. ANa designed and developed notebooks for trap spaces and input propagation and performed verification experiments. SS advised on methodology, supervised and analysed results. ANi and VS prepared figures and wrote the first draft of the manuscript. All authors contributed text and read and approved the final version of the manuscript.

## Supplementary material

All supplementary tables and figures are included in a single pdf file.

## Footnotes

1 https://clinicaltrials.gov/ct2/show/NCT02123264

2 https://www.pharmacodia.com/yaodu/html/v1/chemicals/8651920f3ba2e5f8bcd3e58ba0b48584.html

## Notes

https://gitlab.com/genhotel/rheumatoid-arthritis-large-scale-computational-modeling/-/tree/main

### Competing Interest Statement

The authors have declared no competing interest.

