## Supplementary material for "A large-scale Boolean model of the Rheumatoid Arthritis Fibroblast-Like Synoviocytes predicts drug synergies in the arthritic joint"

**Supplementary Table 1.** The global RA-FLS model comprises 261 nodes. In Table 1, we indicate the biological source of each component.

| No | Global RA-FLS model components | Source (RA-FLS=Rheumatoid Arthritis-Fibroblast-Like Synoviocytes, NR=No Reference, PBMCs= Human Peripheral Blood Mononuclear cells, TH=T Helper cell) |
|---|---|---|
| 1. | ADAMTS4 | RA-FLS |
| 2. | ADAMTS9 | RA-FLS |
| 3. | AKT2 | RA-FLS |
| 4. | APAF1 | RA-FLS |
| 5. | APC | NR |
| 6. | Apoptosis | RA-FLS |
| 7. | ARHGAP35 | NR |
| 8. | ARHGDIB (PBMC) | PBMCs |
| 9. | ARHGEF1 (PBMC) | PBMCs |

| | | |
|---|---|---|
| 10. | ARHGEF2 | RA-FLS |
| 11. | AXIN | NR |
| 12. | BAD | RA-FLS |
| 13. | BAK1 | RA-FLS |
| 14. | BAX | RA-FLS |
| 15. | BCL2 | RA-FLS |
| 16. | BCL2L1 | RA-FLS |
| 17. | BCL2L11 | RA-FLS |
| 18. | BID | RA-FLS |
| 19. | Bone erosion | RA-FLS |
| 20. | BRAF | RA-FLS |
| 21. | CALCINEURIN | RA-FLS |
| 22. | CALM1 | RA-FLS |
| 23. | CALM13 | RA-FLS |
| 24. | CAMK2A | RA-FLS |
| 25. | CASP3 | RA-FLS |
| 26. | CASP8 | RA-FLS |
| 27. | CASP8AP2 | PBMCs |
| 28. | CASP9 | RA-FLS |

| | | |
|---|---|---|
| 29. | CAV1 | RA-FLS |
| 30. | CD14 | RA-FLS |
| 31. | CD19 | PBMCs |
| 32. | CD28 | Synovial tissue, blood, TH1 |
| 33. | CD3E | Synovial tissue, PBMCs, TH1 |
| 34. | CD72 | Blood, synovial fluid |
| 35. | Cell growth/Survival/Proliferation | RA-FLS |
| 36. | CFLAR | RA-FLS |
| 37. | cGAS | RA-FLS |
| 38. | CK1A (CSNK1A1) | RA-FLS |
| 39. | COMMD1 | NR |
| 40. | COMP | RA-FLS |
| 41. | CREB1 | RA-FLS |
| 42. | CRKL | PBMCs, synovial tissue |
| 43. | CSK | RA-FLS |
| 44. | CTNNB | RA-FLS |
| 45. | CTNNB1 | RA-FLS |
| 46. | CTNND1 | Synovial tissue |
| 47. | CXCL10 | RA-FLS |

| | | |
|---|---|---|
| 48. | CXCL11 | RA-FLS |
| 49. | CXCL12 | RA-FLS |
| 50. | CXCL16 | RA-FLS |
| 51. | CXCL8 | RA-FLS |
| 52. | CXCL9 | RA-FLS |
| 53. | CXCR4 | RA-FLS |
| 54. | CXCR6 | RA-FLS |
| 55. | CYBA | RA-FLS |
| 56. | CYBB | NR |
| 57. | CYCS | RA-FLS |
| 58. | DAXX | RA-FLS |
| 59. | DKK1 | RA-FLS |
| 60. | DOCK2 | Synovial tissue |
| 61. | DUSP4 | RA-FLS |
| 62. | DVL1 | RA-FLS |
| 63. | DYNLRB1 | RA-FLS |
| 64. | EGF | RA-FLS |
| 65. | EGFR | RA-FLS |
| 66. | ETS1 | RA-FLS |

| | | |
|---|---|---|
| 67. | FADD | RA-FLS |
| 68. | FAS | RA-FLS |
| 69. | FASLG | RA-FLS |
| 70. | FGF1 | RA-FLS |
| 71. | FGFR1 | RA-FLS |
| 72. | FN1 | RA-FLS |
| 73. | FOS | RA-FLS |
| 74. | FOXO1 | RA-FLS |
| 75. | FOXO3 | RA-FLS |
| 76. | FRIZZLED | PBMCs, synovial tissue |
| 77. | FYN | Macrophages, synovial tissue |
| 78. | GAB1 | RA-FLS |
| 79. | GAB2 | RA-FLS |
| 80. | GNAI3 | PBMCs |
| 81. | GNB1 | PBMCs |
| 82. | GRB2 | RA-FLS |
| 83. | GSK3b | RA-FLS |
| 84. | HCST | PBMCs, synovial tissue |
| 85. | ICAM1 | RA-FLS |

| | | |
|---|---|---|
| 86. | ICAM2 | RA-FLS |
| 87. | ICAM3 | RA-FLS |
| 88. | IFNA1 | NR |
| 89. | IFNB1 | RA-FLS |
| 90. | IFNAR1 | RA-FLS |
| 91. | IFNAR2 | Macrophages |
| 92. | IGF1 | RA-FLS |
| 93. | IGF1R | RA-FLS |
| 94. | IKBA | RA-FLS |
| 95. | IKBKE | RA-FLS |
| 96. | IKK | RA-FLS |
| 97. | IL17A | RA-FLS |
| 98. | IL17RA | RA-FLS |
| 99. | IL18 | RA-FLS |
| 100 | IL1A | RA-FLS |
| 101 | IL1B | RA-FLS |
| 102 | IL1R1 | RA-FLS |
| 103 | IL6 | RA-FLS |
| 104 | IL6R | RA-FLS |

| | | |
|---|---|---|
| 105 | IL6ST | RA-FLS |
| 106 | IL7 | RA-FLS |
| 107 | Inflammation | RA-FLS |
| 108 | ILK | PBMCs, TH1 |
| 109 | IRAK1 | RA-FLS |
| 110 | IRAK4 | RA-FLS |
| 111 | IRF1 | RA-FLS |
| 112 | IRF3 | RA-FLS |
| 113 | IRF5 | RA-FLS |
| 114 | IRF7 | Chondrocytes, TH1, macrophages |
| 115 | IRF8 | NR |
| 116 | IRF9 | RA-FLS |
| 117 | ISGF3 | RA-FLS |
| 118 | ITAM | Synovial tissue |
| 119 | ITBG2 | RA-FLS |
| 120 | ITGAL | RA-FLS |
| 121 | ITGAV | RA-FLS |
| 122 | JAK1 | RA-FLS |
| 123 | JAK2 | RA-FLS |

| | | |
|---|---|---|
| 124 | JAK3 | RA-FLS |
| 125 | JUN | RA-FLS |
| 126 | KLRK1 | PBMCs |
| 127 | LBP | Synovial tissue, synovial fluid |
| 128 | LCK | Synovial tissue |
| 129 | LCP2 | Synovial tissue, PBMCs, TH1 |
| 130 | LEFTY2 | RA-FLS |
| 131 | LRP156 | Synovial tissue |
| 132 | LTBP1 | RA-FLS |
| 133 | LY96 | PBMCs |
| 134 | MAP2K1 | RA-FLS |
| 135 | MAP2K2 | RA-FLS |
| 136 | MAP2K3 | RA-FLS |
| 137 | MAP2K4 | RA-FLS |
| 138 | MAP2K6 | RA-FLS |
| 139 | MAP2K7 | RA-FLS |
| 140 | MAP3K1 | RA-FLS |
| 141 | MAP3K14 | RA-FLS |
| 142 | MAP3K234 | RA-FLS |

| | | |
|---|---|---|
| 143 | MAP3K5 | RA-FLS |
| 144 | MAP3K7 | RA-FLS |
| 145 | MAP3K8 | RA-FLS |
| 146 | MAP4K1 | Synovial tissue |
| 147 | MAP4K4 | RA-FLS |
| 148 | MAPK1 | RA-FLS |
| 149 | MAPK14 | RA-FLS |
| 150 | MAPK3 | RA-FLS |
| 151 | MAPK7 | Synovial tissue |
| 152 | MAPK8 | RA-FLS |
| 153 | MAPK9 | RA-FLS |
| 154 | MAPKAPK2 | RA-FLS |
| 155 | Matrix degradation | RA-FLS |
| 156 | MCL1 | RA-FLS |
| 157 | MDM2 | RA-FLS |
| 158 | MICB | PBMCs |
| 159 | MMP1 | RA-FLS |
| 160 | MMP13 | RA-FLS |
| 161 | MMP3 | RA-FLS |

| | | |
|---|---|---|
| 162 | MMP9 | RA-FLS |
| 163 | MTORC1 | RA-FLS |
| 164 | MYD88 | RA-FLS |
| 165 | NCF | RA-FLS |
| 166 | NECTIN1 | Synovial tissue |
| 167 | NECTIN3 | NR |
| 168 | NFAT5 | RA-FLS |
| 169 | NFATC1 | Synovial tissue |
| 170 | NFKB | RA-FLS |
| 171 | NFKB1 | RA-FLS |
| 172 | NLK | Macrophages and blood |
| 173 | NRAS | RA-FLS |
| 174 | NRF1 | Synovial tissue |
| 175 | OSCAR | Synovial tissue, blood |
| 176 | p38MAPK | RA-FLS |
| 177 | PDE3A | NR |
| 178 | PDGFA | RA-FLS |
| 179 | PDGFRA | RA-FLS |
| 180 | PIK3R5 | RA-FLS |

| | | |
|---|---|---|
| 181 | PLCG1 | TH1 |
| 182 | PLXNB1 | RA-FLS |
| 183 | PMAIP1 | RA-FLS |
| 184 | PP2A | RA-FLS |
| 185 | PPP3CB | Synovial tissue, TH1 |
| 186 | PRKACG | PBMCs |
| 187 | PRKCD | Synovial tissue, macrophages |
| 188 | PRKCH | PBMCs |
| 189 | PRRT2 | NR |
| 190 | PTEN | RA-FLS |
| 191 | PTK2B | RA-FLS |
| 192 | PTPN11 | RA-FLS |
| 193 | PTPN6 | Synovial tissue, macrophages |
| 194 | PTPRC | RA-FLS |
| 195 | RAB5A | RA-FLS |
| 196 | RAC12 | RA-FLS |
| 197 | RAF1 | RA-FLS |
| 198 | RANK | RA-FLS |
| 199 | RANKL | RA-FLS |

| | | |
|---|---|---|
| 200 | RAP1B | RA-FLS |
| 201 | RAPGEF1 | PBMCs, synovial fluid |
| 202 | RASGRP3 | PBMCs, macrophages, synovial fluid |
| 203 | RELA | RA-FLS |
| 204 | RHOA | RA-FLS |
| 205 | RIPK1 | PBMCs, TH1 |
| 206 | RPS6KA1 | Synovial tissue, PBMCs, TH1) |
| 207 | RPS6KA5 | Synovial tissue |
| 208 | RPS6KB1 | RA-FLS |
| 209 | SARM1 | RA-FLS |
| 210 | SFRP5 | RA-FLS |
| 211 | SH2D1A | RA-FLS |
| 212 | SHC2 | RA-FLS |
| 213 | SHP2 | RA-FLS |
| 214 | SOCS3 | RA-FLS |
| 215 | SOS1 | RA-FLS |
| 216 | SRC | RA-FLS |
| 217 | SRF | PBMCs |
| 218 | STAT1 | RA-FLS |

| | | |
|---|---|---|
| 219 | STAT2 | RA-FLS |
| 220 | STAT3 | RA-FLS |
| 221 | SYK | NR |
| 222 | TAB1 | RA-FLS |
| 223 | TAB2 | NR |
| 224 | TAK1 | RA-FLS |
| 225 | TBK1 | RA-FLS |
| 226 | LEF | Macrophages |
| 227 | TCF | RA-FLS |
| 228 | TGFB1 | RA-FLS |
| 229 | TGFBR1 | RA-FLS |
| 230 | TICAM1 | RA-FLS |
| 231 | TICAM2 | NR |
| 232 | TIRAP | RA-FLS |
| 233 | TLR2 | RA-FLS |
| 234 | TLR4 | RA-FLS |
| 235 | TLR5 | RA-FLS |
| 236 | TNF | RA-FLS |
| 237 | TNFAIP3 | Synovial tissue, PBMCs, macrophages |

| | | |
|---|---|---|
| 238 | TNFRSF10A | RA-FLS |
| 239 | TNFRSF10B | RA-FLS |
| 240 | TNFRSF1A | RA-FLS |
| 241 | TNFRSF1B | RA-FLS |
| 242 | TNFSF11 | RA-FLS |
| 243 | TP53 | RA-FLS |
| 244 | TP73 | RA-FLS |
| 245 | TRADD | PBMCs, macrophages |
| 246 | TRAF2 | RA-FLS |
| 247 | TRAF3 | RA-FLS |
| 248 | TRAF3IP2 | RA-FLS |
| 249 | TRAF5 | RA-FLS |
| 250 | TRAF6 | RA-FLS |
| 251 | TXK | RA-FLS |
| 252 | TYK2 | Blood |
| 253 | TYROBP | Synovial tissue |
| 254 | VAV1 | Synovial tissue |
| 255 | VAV2 | Synovial tissue |
| 256 | WNT | Synovial tissue |

| | | |
|---|---|---|
| 257 | WNT5A | RA-FLS |
| 258 | YWHAQ | RA-FLS |
| 259 | YY1 | RA-FLS |
| 260 | ZAP70 | PBMCs, TH1) |
| 261 | ZC3H12A | RA-FLS |

**Supplementary Table 2.** Biological scenarios tested using the unmodified and modified RA-FLS global models and the phenotype-specific modules.

† displays RA-FLS specific scenarios reproduced with additional conditions of (* known biological behaviour, ** model conditions added to reproduce the RA-FLS biological scenarios. The added model conditions are not experimentally proven).

| **Phenotypes** | **Biological scenarios** | **References** | **Validation using the phenotype-specific module** | **Validation using the RA-FLS global model** | **Validation using the modified global RA-FLS model** |
|---|---|---|---|---|---|
| **Inflammation** | 1) Anti-TNF-α monoclonal antibody results in a decrease in synovial inflammation<br>**TNF KO** | Matsuno et al., 2002 | Yes | Yes | Yes |
| | 2) Synovial inflammation was significantly exacerbated by TNF-α administration<br>**TNF ON** | Matsuno et al., 2002 | Yes<br>(by manually activating NFKB, which is a ***known** biological condition regulating TNF) | Yes<br>(by manually activating NFKB, which is a ***known** biological condition regulating TNF | Yes<br>(by manually activating NFKB, which is a ***known** biological condition regulating TNF |
| | 3) Interleukin 6 knockout mice are resistant to antigen-induced experimental | Boe et al., 1999 | Yes | Yes | Yes |

| | | | | | |
|---|---|---|---|---|---|
| | arthritis<br>**IL6 KO** | | | | |
| | 4) Anti-IL17A antibody reduces joint inflammation in collagen-induced arthritis (CIA)<br>**IL17 KO** | Lubberts et al., 2004; Nakae et al.,2003 | Yes | Yes | Yes |
| | 5) Secreted frizzled-related protein 5 suppresses inflammatory response in rheumatoid arthritis fibroblast-like synoviocytes<br>**SFRP5 ON** | Kwon et al., 2014 | Yes | Yes | Yes |
| **Bone erosion** | 1) Secreted frizzled-related protein 5 suppresses inflammatory response in rheumatoid arthritis fibroblast-like synoviocytes<br>**SFRP5 ON** | Kwon et al., 2014 | Yes | Yes | Yes |
| | 2) RANKL (TNFSF11) expressed on synovial fibroblasts is involved in rheumatoid bone destruction by inducing osteoclastogenesis<br>**TNFSF11 ON** | Takayanagi et al., 2000 | Yes | Yes | Yes |
| | 3) RANKL (TNFSF11) knockout mice are protected from bone erosion in a serum transfer model of arthritis<br>**TNFSF11 KO** | Pettit et al., 2001 | Yes<br>(by manually activating SFRP5, a **model condition) | Yes<br>(by manually activating SFRP5, a **model condition) | Yes<br>(by manually activating SFRP5, a **model condition) |
| | 4) Wnt5a cKO mice | MacLauchla | Yes | Yes | Yes |

| | | | | | |
|---|---|---|---|---|---|
| | were resistant to arthritis development **WNT5A KO** | n et al., 2017 | (by manually activating SFRP5, a **model condition) | (by manually activating SFRP5, a **model condition) | (by manually activating SFRP5, a **model condition) |
| | 5) Fas receptor induces apoptosis of synovial bone and cartilage progenitor populations and promotes bone loss in antigen-induced arthritis **FASFASLG ON** | Li et al., 2017; Lazić Mosler et al., 2019 | Yes | Yes | Yes |
| **Cell proliferation** | 1) PDGF stimulates the proliferation of synovial fibroblasts **PDGFA ON** | Sandler et al., 2006; Kameda et al., 2006 | Yes | Yes | Yes |
| | 2) PDGF stimulates the growth of synovial fibroblast via interference with the Akt signalling pathway **PDGFA ON_AKT2 ON** | Terabe et al., 2009 | Yes | Yes | Yes |
| | 3) Downregulation of FoxO1 is required to promote the survival of fibroblast-like synoviocytes in rheumatoid arthritis **FOXO1 OFF** | Grabiec et al., 2015 | Yes | Yes | Yes |
| | 4) TGFbeta exerts its growth and antiapoptotic effects on fibroblasts by activation of the PI3Kinase-AKT pathway **TGFB1 ON_PIK3R5 ON** | Kim G et al., 2002 | Yes | Yes | Yes |

| | | | | | |
|---|---|---|---|---|---|
| **Matrix degradation** | 1) Inhibition of MMP1 alone results in a significant reduction of cartilage invasion by RASFs **MMP1 OFF** | Rutkauskaite et al., 2004 | Yes | Yes | Yes |
| | 2) MMP-9 stimulates RA synovial fibroblast-mediated inflammation and degradation of cartilage **MMP9 ON** | Xue et al., 2014 | Yes | Yes | Yes |
| **Apoptosis** | 1) Fas receptor induces apoptosis of synovial bone and cartilage progenitor populations and promotes bone loss in antigen-induced arthritis **FASFASLG ON -** original model, **FASFASLG ON_MIR192 ON (modified)** - modified_model | Lazić Mosler et al., 2019 | Original module - Yes, Modified module - Yes (by manually activating MIR192_rna to deactivate the dominant CAV1 negative regulator, a **model condition) | Yes | Yes (by manually activating MIR192_rna to deactivate the dominant CAV1 negative regulator, a **model condition) |
| | 2) TNFα can induce apoptosis in RA-FLS when NF-κB is inhibited **TNF ON_NFKB OFF-** original model, **TNF ON_NFKB OFF_MIR192 ON (modified)** - modified model | Zhang et al., 2000 | Original module - Yes, Modified module - Yes (by manually activating MIR192_rna to deactivate the dominant CAV1 negative regulator, a **model condition) | Yes | Yes (by manually activating MIR192_rna to deactivate the dominant CAV1 negative regulator, a **model condition) |
| | 3) In RA-FLS, phosphorylation of AKT protects against Fas-induced apoptosis through inhibition of Bid cleavage. | García et al., 2010 | Original module - No, Modified module - Yes | No | Yes |

| | | | | | |
|---|---|---|---|---|---|
| | **AKT2 ON_FASLG ON_BID OFF** | | | | |
| | 4) TGFbeta exerts its growth and antiapoptotic effects on fibroblasts by activation of the PI3Kinase-AKT pathway **TGFB1 ON_PIK3R5 ON** | Kim G et al., 2002 | Original module - unfixed, Modified module - Yes | Unfixed | Yes |

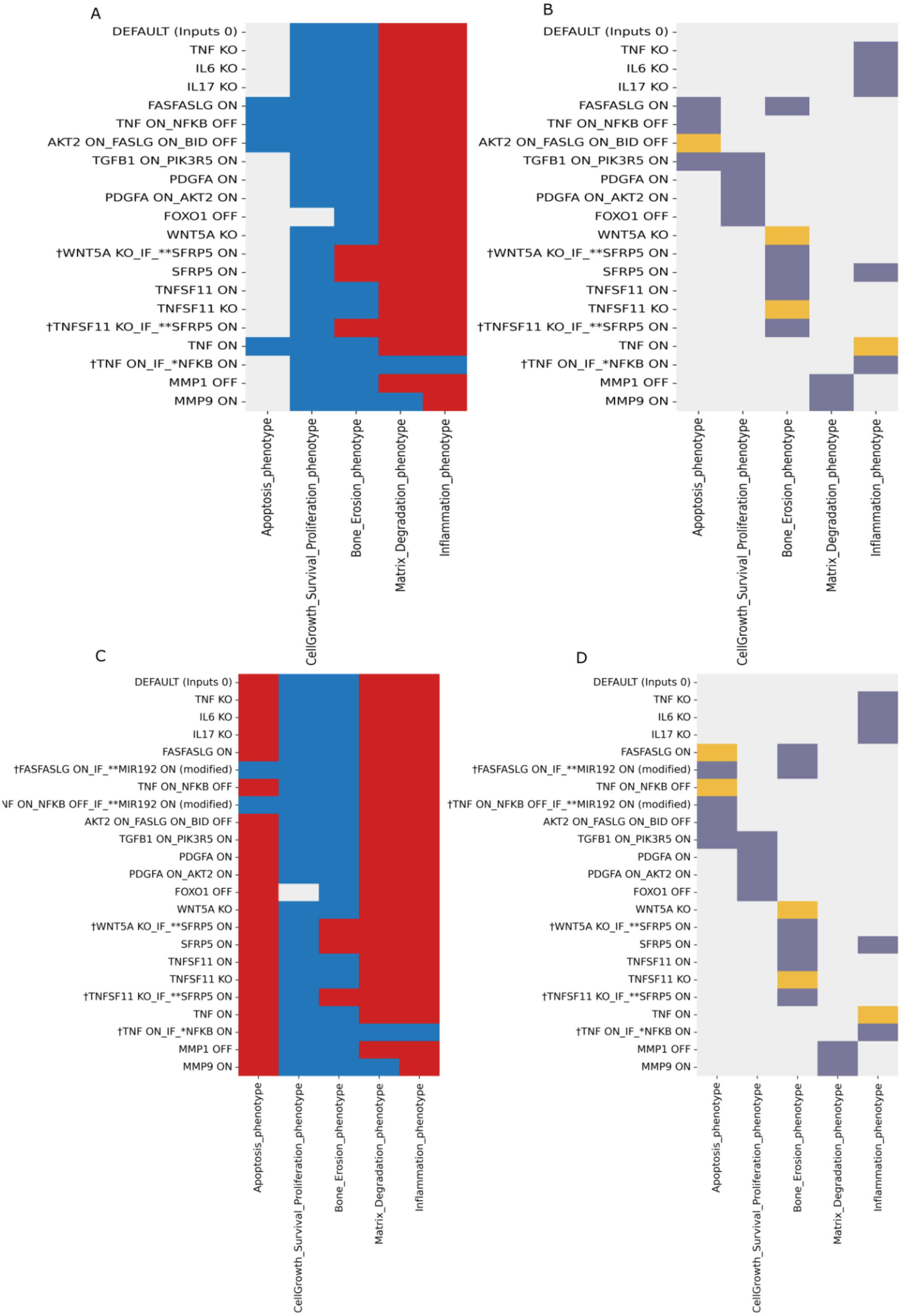
A
DEFAULT (Inputs 0)
TNF KO
IL6 KO
IL17 KO
FASFASLG ON
TNF ON_NFKB OFF
AKT2 ON_FASLG ON_BID OFF
TGFB1 ON_PIK3R5 ON
PDGFA ON
PDGFA ON_AKT2 ON
FOXO1 OFF
WNT5A KO
†WNT5A KO_IF_**SFRP5 ON
SFRP5 ON
TNFSF11 ON
TNFSF11 KO
†TNFSF11 KO_IF_**SFRP5 ON
TNF ON
†TNF ON_IF_*NFKB ON
MMP1 OFF
MMP9 ON
Apoptosis_phenotype
CellGrowth_Survival_Proliferation_phenotype
Bone_Erosion_phenotype
Matrix_Degradation_phenotype
Inflammation_phenotype
B
DEFAULT (Inputs 0)
TNF KO
IL6 KO
IL17 KO
FASFASLG ON
TNF ON_NFKB OFF
AKT2 ON_FASLG ON_BID OFF
TGFB1 ON_PIK3R5 ON
PDGFA ON
PDGFA ON_AKT2 ON
FOXO1 OFF
WNT5A KO
†WNT5A KO_IF_**SFRP5 ON
SFRP5 ON
TNFSF11 ON
TNFSF11 KO
†TNFSF11 KO_IF_**SFRP5 ON
TNF ON
†TNF ON_IF_*NFKB ON
MMP1 OFF
MMP9 ON
Apoptosis_phenotype
CellGrowth_Survival_Proliferation_phenotype
Bone_Erosion_phenotype
Matrix_Degradation_phenotype
Inflammation_phenotype
C
DEFAULT (Inputs 0)
TNF KO
IL6 KO
IL17 KO
FASFASLG ON
†FASFASLG ON_IF_**MIR192 ON (modified)
TNF ON_NFKB OFF
NF ON_NFKB OFF_IF_**MIR192 ON (modified)
AKT2 ON_FASLG ON_BID OFF
TGFB1 ON_PIK3R5 ON
PDGFA ON
PDGFA ON_AKT2 ON
FOXO1 OFF
WNT5A KO
†WNT5A KO_IF_**SFRP5 ON
SFRP5 ON
TNFSF11 ON
TNFSF11 KO
†TNFSF11 KO_IF_**SFRP5 ON
TNF ON
†TNF ON_IF_*NFKB ON
MMP1 OFF
MMP9 ON
Apoptosis_phenotype
CellGrowth_Survival_Proliferation_phenotype
Bone_Erosion_phenotype
Matrix_Degradation_phenotype
Inflammation_phenotype
D
DEFAULT (Inputs 0)
TNF KO
IL6 KO
IL17 KO
FASFASLG ON
†FASFASLG ON_IF_**MIR192 ON (modified)
TNF ON_NFKB OFF
†TNF ON_NFKB OFF_IF_**MIR192 ON (modified)
AKT2 ON_FASLG ON_BID OFF
TGFB1 ON_PIK3R5 ON
PDGFA ON
PDGFA ON_AKT2 ON
FOXO1 OFF
WNT5A KO
†WNT5A KO_IF_**SFRP5 ON
SFRP5 ON
TNFSF11 ON
TNFSF11 KO
†TNFSF11 KO_IF_**SFRP5 ON
TNF ON
†TNF ON_IF_*NFKB ON
MMP1 OFF
MMP9 ON
Apoptosis_phenotype
CellGrowth_Survival_Proliferation_phenotype
Bone_Erosion_phenotype
Matrix_Degradation_phenotype
Inflammation_phenotype

**Supplementary Figure 1.** Heatmap displaying the trap spaces for the tested biological scenarios in the unmodified global model (A), modified global model (C) and the comparison between expected and obtained values shown with colour codes for the unmodified (B) and the modified (D) models with DEFAULT inputs set to zero. The y-axis shows all the tested scenarios' names, as mentioned in table 2, in regard to all the five phenotypes as outcomes on the X-axis. † represent scenarios where additional conditions were given as a known biological behaviour* or as model conditions**.

**Trap spaces colour codes:** -1 (unfixed), 0 (OFF), 1 (ON)

**Expectation graph colour codes:**

**Score: expected value, obtained value; 1**: Yes [OFF, OFF & ON, ON]; **0**: No [ON, OFF & OFF, ON (conflict)], **-1**: Undefined

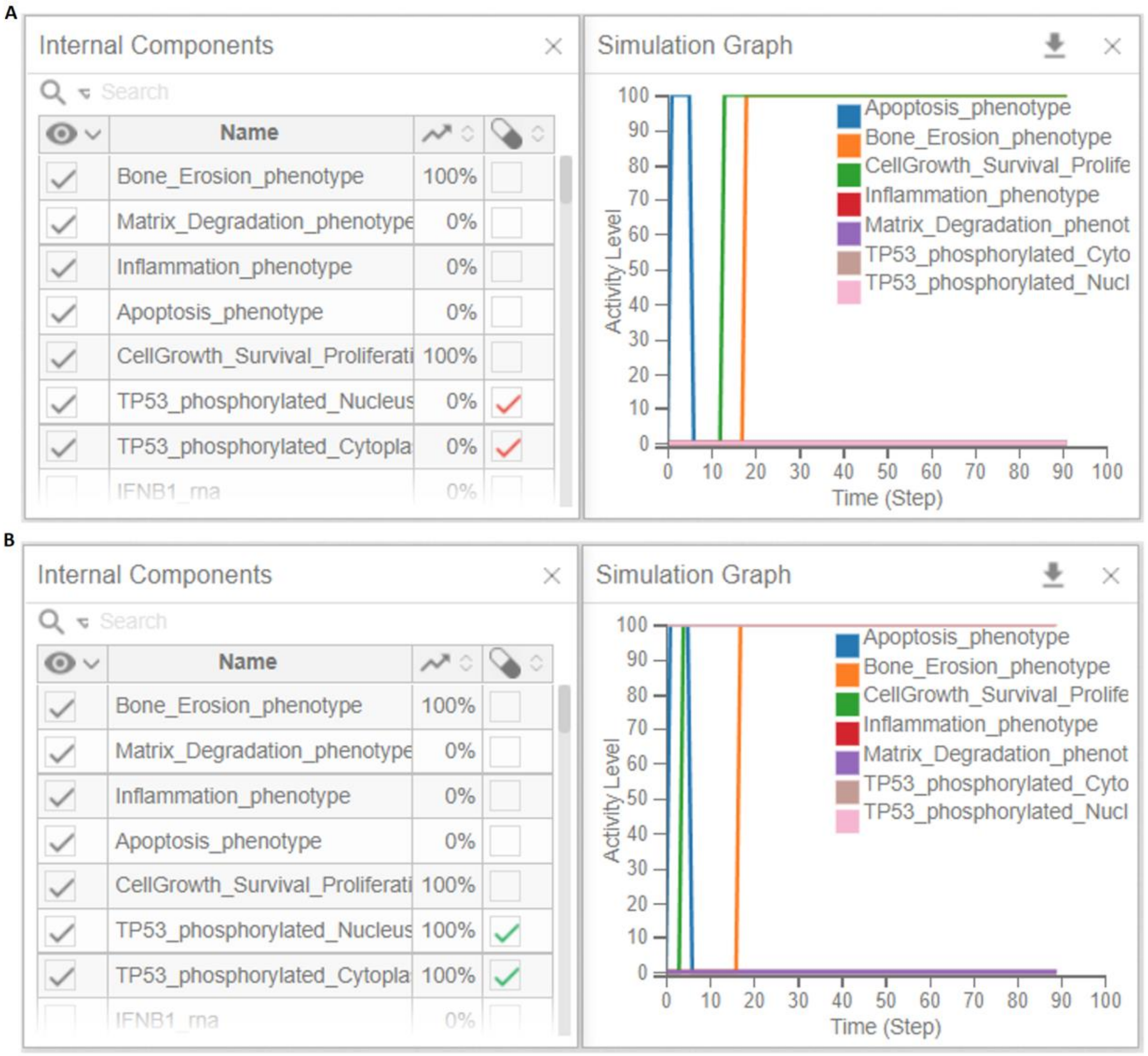


**Supplementary Figure 2**. Simulation results using the Cell Collective platform with the modified RA-FLS model. All inputs are kept inactive, and two different conditions of TP53 are tested, OFF (A) and ON (B). When TP53 is kept ON, apoptosis, matrix degradation and inflammation remain OFF, while bone erosion and cell proliferation remain ON (A). However, turning TP53 OFF under the same conditions did not impact the phenotypic outcomes (B).

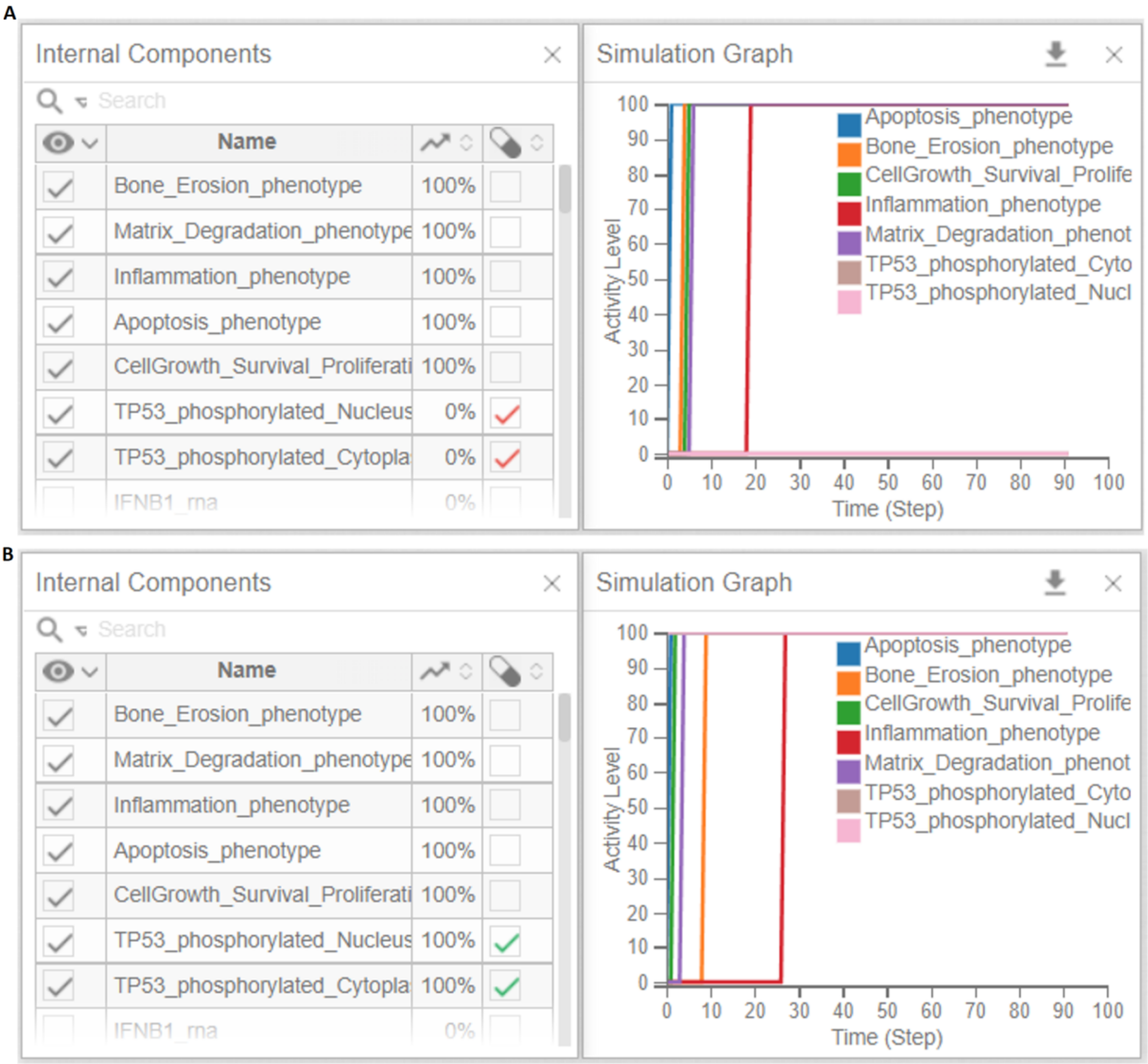


**Supplementary Figure 3**. Simulation results using the Cell Collective platform with the modified RA-FLS model. All inputs are kept active, and two different conditions of TP53 are tested, OFF (A) and ON (B). All phenotypes remain active in both tested conditions.

**Supplementary Table 3. Direct upstream regulators of the five phenotypes and available drugs that target them**

* drugs used in RA

| Phenotypes | Logical formulae | Direct upstream regulators | Drugs targeting direct upstream regulators | No of clinical trials for the drug | In clinical trials/*in vitro* /*in vivo* |
|---|---|---|---|---|---|
| Apoptosis | !CAV1_rna&(CASP3_phosphorylated\|CASP8\|TNFRSF10A_rna\|TNFRSF10B_rna | CAV1<br>CASP3<br>CASP8<br>TNFRSF10A<br>TNFRSF10B | **CAV1: Pamidronate**<br><br>**Zoledronic acid***<br><br>**Incadronate** | **56**<br><br>**380** | **Pamidronate**<br>Phase1/2/3/4 myeloma, osteoporosis, arthroplasty, osteopenia, cancers (https://clinicaltrials.gov/ct2/results?cond=&term=Pamidronate&cntry=&state=&city=&dist=**)**<br><br>**Zoledronic acid***<br>Phase1/2/3/4<br>RA, osteoporosis, osteopenia, cancers, Parkinson's Disease, multiple myeloma, (https://clinicaltrials.gov/ct2/results?term=Zoledronic+acid&age_v=&gndr=&type=&rslt=&Search=Apply)<br><br>**Incadronate**<br>RA (Zhao et al. 2006)<br>prostate cancer (Iguchi et al. 2006) |
| Cell proliferation | NFKB/N_complex\|YWHAQ_rna\|CREB1_phosphorylated\|BCL2_phosphorylated\|RPS6KB1_phosphorylated\|TNF_Secreted_space_Molecules\|ADAMTS9_rna\|p38MAPK_phosphorylated | TNF<br>NFKB_complex<br>CREB1<br>BCL2<br>RPS6KB1<br>ADAMTS9<br>P38MAPK<br>YWHAQ** | **NFKB: Bortezomib***<br><br>**Salvianolic acid B*** (SA-B)<br><br>**P38MAPK: CDD-450*** , | **1040**<br><br>**2** | **Bortezomib**<br>Phase2/3/4<br>RA, multiple myeloma, lymphoma, leukaemia, different cancers<br>(https://clinicaltrials.gov/ct2/results?cond=&term=Bortezomib&cntry=&state=&city=&dist=), (Chen et al. 2011)<br><br>Rat model (Xia et al. 2018)<br><br>**CDD-450**<br>Phase 2<br>RA, psoriatic arthritis<br>**(**https://clinicaltrials.gov/ct2/results?cond=&term=CDD-450&cntry=&state=&city=&dist=**),**<br>(https://doi.org/10.1038/s41584-018-0003-y) |

| | | | | | |
|---|---|---|---|---|---|
| | | | **SD-0006** | **16** | **SD-0006**<br>Cancers, type 2 diabetes, Alzheimer's Disease<br>Phase2/3/4<br>**(**https://clinicaltrials.gov/ct2/results?term=SD+0006&rank=1#rowId**0)** |
| | | | **R-1487 *** | | Phase 1 (Goldstein et al. 2011) |
| | | | **BMS-582949** | **4** | **BMS-582949**<br>Phase 1/2<br>RA, psoriasis, atherosclerosis<br>(https://clinicaltrials.gov/ct2/results?cond=&term=BMS-582949&cntry=&state=&city=&dist=) |
| | | | **Dilmapimod*** | **10** | **Dilmapimod**<br>phase 1/2 RA, pulmonary Disease<br>(https://clinicaltrials.gov/ct2/results?term=Dilmapimod&draw=2&rank=2#rowId1)<br>**Discontinued** (Lakshmi et al. 2017) |
| | | | **Doramapimod*** | **11** | **Doramapimod**<br>Phase 2<br>RA, psoriasis, Crohn's Disease<br>(https://clinicaltrials.gov/ct2/results?term=Doramapimod&draw=2&rank=2#rowId1) |
| | | | **Losmapimod** | **15** | **Losmapimod**<br>Phase2, Facioscapulohumeral Muscular Dystrophy (FSHD), glomerulosclerosis, focal segmental, pulmonary disease, chronic obstructive, acute coronary syndrome, atherosclerosis<br>(https://clinicaltrials.gov/ct2/results?cond=&term=Losmapimod&cntry=&state=&city=&dist=) |
| | | | **LY2228820** | **5** | **LY2228820**<br>Phase 1, metastatic breast cancer, adult glioblastoma, epithelial ovarian cancer, advanced cancer<br>(https://clinicaltrials.gov/ct2/results?cond=&term=LY2228820&cntry=&state=&city=&dist=) |

| | | | | | |
|---|---|---|---|---|---|
| | | | **PH-797804 *** | **16** | **PH-797804**<br>Phase 2, RA, chronic obstructive pulmonary disease, pulmonary disease, chronic obstructive, osteoarthritis, neuralgia, postherpetic neuralgia (https://clinicaltrials.gov/ct2/results?cond=&term=PH-797804&cntry=&state=&city=&dist=) |
| | | | **VX-745 *** | **5** | **VX-745**<br>Phase 2, Alzheimer's Disease, Huntington's Disease (https://clinicaltrials.gov/ct2/results?cond=&term=VX-745&cntry=&state=&city=&dist=), RA(Haddad 2001) |
| | | | **BCL2: Obatoclax** | **20** | **Obatoclax**<br>Phase 1/2 of different leukaemia and lymphoma (https://clinicaltrials.gov/ct2/results?cond=&term=Obatoclax&cntry=&state=&city=&dist=), (Or et al. 2020) |
| | | | **CREB1: 666-15** | | **666-15**<br>*In vivo* mice studies (Xie et al. 2019; Li et al. 2016) |
| | | | **YWHAQ** targeted by upstream TF **FOXO1: AS1842856** | | **AS1842856**<br>(Zou et al. 2014), *In vivo* mice study (Nagashima et al. 2010), review (Calissi et al. 2021) |
| Inflammation | CXCL11\|IRF7\|CXCL8\|IRF1\|IFNB1_rna\|IL1B_Secreted_space_Molecules\|CXCL9\|TNF_Secreted_space_Molecules\|IL6_Secreted_space_Molecules\|NFKB/N_complex\|IL17A_Secreted_space_M | TNF<br>IL6<br>IL17A<br>IL1B<br>NFKB<br>CXC11<br>IRF7<br>CXCL8<br>IFNB1<br>CXCL9<br>CXCL10<br>IFNA1<br>IRF5<br>IRF1 | **CXCL8: Carboxyamidotriazole**<br><br>**Rivanicline**<br><br>**CXCL10: BMS-936557** | **10**<br><br><br><br>**2** | **Carboxyamidotriazole**<br>Phase1/2 of different cancers (https://clinicaltrials.gov/ct2/results?cond=&term=Carboxyamidotriazole&cntry=&state=&city=&dist=)<br><br>*In vitro* study (Spoettl et al. 2007)<br><br>**BMS-936557**<br>Phase2 colitis, ulcerative (Mayer et al. 2014), Crohn's Disease (https://clinicaltrials.gov/ct2/results?term=BMS-936557&draw=2&rank=1#rowId0) |

| | | | | | |
|---|---|---|---|---|---|
| | olecules\|CXCL10\|IFNA1_rna\|IRF5_Secreted_space_Molecules | | **CXCL11:** targeting by upstream IRF3 **Piceatannol**<br><br>**ADU-S100**<br><br>**IFNA1: Interferon alfa-n3**<br><br>**IFNB1: PEGylated IFN beta 1-a** | 1<br><br>3<br><br>11<br><br>15 | **Piceatannol** Preclinical (https://clinicaltrials.gov/ct2/show/NCT04983017?term=Piceatannol&draw=2&rank=1), In vivo mice study (Dang et al. 2004)<br><br>**ADU-S100** Phase1/2 of cancer/lymphoma (https://clinicaltrials.gov/ct2/results?cond=&term=ADU-S100&cntry=&state=&city=&dist=)<br><br>**Interferon alfa-n3** **(**https://go.drugbank.com/drugs/DB05258), Phase1/2 HIV infections, different cancer/lymphoma/myeloma (https://clinicaltrials.gov/ct2/results?cond=&term=Interferon+alfa-n3&cntry=&state=&city=&dist=)<br><br>**PEGylated IFN beta 1-a** Phase3 multiple sclerosis, SARS-CoV Infection (https://clinicaltrials.gov/ct2/results?cond=&term=PEGylated+IFN+beta+1-a&cntry=&state=&city=&dist=), (Calabresi et al. 2014) |
| Bone erosion | Osteoclastogenesis - IL17A_Secreted_space_Molecules\|TNFSF11_rna\|IL1B_Secreted_space_Molecules\|TNF_Secreted_space_Molecules\|FOS\|JUN_phosphorylated\|IL7\|NFATC1_Secreted_sp | TNF IL17A IL1B IL7 TNFRSF11 FOS JUN NFATC1 | **JUN, FOS (AP-1)** **T-5224,**<br><br>**Acitretin**<br><br>**IL7*** | <br><br>44<br><br>186 | **T-5224** *In vivo* mice study (Ishida et al. 2015)<br><br>**Acitretin** Phase4 psoriasis, Phase1/2 melanoma, HIV infections, Alzheimer's Disease (https://clinicaltrials.gov/ct2/results?cond=&term=acitretin&cntry=&state=&city=&dist=)<br><br>**Certolizumab pegol** |

| | | | | | |
|---|---|---|---|---|---|
| | ace_Molecules | | **Certolizumab pegol**<br><br>**GSK2618960**<br><br>**CYT107**<br><br>**TNFRSF11* Denosumab**<br><br>**NFATC1** | <br><br>**3**<br><br>**259**<br><br>**81** | RA, psoriasis, colitis, spondylitis, Crohn's Disease (https://clinicaltrials.gov/ct2/results?cond=&term=Certolizumab+pegol&cntry=&state=&city=&dist=)<br><br>**GSK2618960**<br>Multiple sclerosis<br>**(https://clinicaltrials.gov/ct2/show/NCT01808482)**<br><br>**CYT107**<br>Cancer, HIV, autoimmune disease (https://clinicaltrials.gov/ct2/results?term=il7&draw=2&rank=18#rowId17)<br><br>**Denosumab**<br>MOA (Bruhn 2010), Phase2/3/4 RA, osteoporosis, cystic fibrosis, bone resorption, breast cancer, cancers (https://clinicaltrials.gov/ct2/results?term=Denosumab&age_v=&gndr=&type=&rslt=&phase=1&Search=Apply)<br><br>**INCA-6**<br>*In vitro* and cells (Roehrl et al. 2004), Preclinical |
| Matrix degradation | MMP13\|MMP9\|MMP1\|ADAMTS4\|MMP3 | MMP13<br>MMP9<br>MMP1<br>MMP3<br>ADAMTS4 | MMP9, MMP3, MMP13,MMP1<br>**CTS-1027**<br><br>**Marimastat** | **4**<br><br>**4** | **CTS-1027**<br>**Hepatitis**<br>(https://clinicaltrials.gov/ct2/show/NCT00925990), https://drugs.ncats.io/drug/2QD3F58224<br>**It was discontinued due to laboratory abnormalities and adverse events in a subset of clinical trial participants.**<br><br>**Marimastat**<br>Glioblastoma, breast, ovarian and small and non-small cell lung cancer (https://clinicaltrials.gov/ct2/results?cond=&term=Marimastat+&cntry=&state=&city=&dist=)<br>**It was discontinued as it failed to show superior efficacy over either standard chemotherapy or placebo.** |

| | | | | | |
|---|---|---|---|---|---|
| | | | **Cipemastat***<br><br>**Apratastat (TMI-005)** *<br><br>**MMP9 Andecaliximab** (GS-5745)<br><br>**Salvianolic acid B*** (SA-B)<br><br>**Curcumin*** | **13**<br><br>**1**<br><br>**2**<br><br>**3** | **Cipemastat**<br>Inhibits MMP-1, MMP-3 and MMP-9 but did not prevent the progression of joint damage in RA **(**Vandenbroucke, R., Libert, C, 2014)<br>**It was discontinued as it failed to improve radiographic scores**<br><br>**Apratastat**<br>TACE and MMPs inhibitor (Thabet and Huizinga 2006)<br>**It was discontinued due to a lack of efficacy**<br><br>**Andecaliximab** (GS-5745)<br>Phase 1 gastroesophageal junction adenocarcinoma (2022) (Yoshikawa et al. 2022), Phase1 RA (Gossage et al. 2018), Phase3 gastric adenocarcinoma (https://clinicaltrials.gov/ct2/show/NCT02545504?term=Andecaliximab&phase=2&draw=2&rank=1), phase2/3 ulcerative colitis (https://clinicaltrials.gov/ct2/show/NCT02520284?term=Andecaliximab&phase=2&draw=2&rank=2)<br><br>**Salvianolic acid B*** (SA-B)<br>Preclinical fatty liver disease(https://clinicaltrials.gov/ct2/show/NCT05076058?term=Salvianolic+acid+B&draw=2&rank=2),<br>Hepatitis B (Liu et al. 2002), in vitro cells, and in vivo mice (Lin et al. 2007)<br><br>**Curcumin***<br>Phase 1/2<br>RA, type 2 diabetes<br>(https://clinicaltrials.gov/ct2/show/NCT01052597),<br>(https://clinicaltrials.gov/ct2/show/NCT00752154?term=curcumin&cond=Rheumatoid+Arthritis&draw=2&rank=1)*,<br>(Dai et al. 2018),(Chuengsamarn et al. |

| | | | | | |
|---|---|---|---|---|---|
| | | | | | 2014) |
| | | | **MMP3: Prinomastat** | | **Prinomastat** Phase 3 lung and prostate cancer, brain and central nervous system tumours (https://clinicaltrials.gov/ct2/results?cond=&term=Prinomastat&cntry=&state=&city=&dist=) |
| | | | **Batimastat** | | **Batimastat** (Kumar et al. 2010) Duchenne muscular dystrophy (DMD), Internal resorption (IR), Tuberculosis (TB) (Tang et al. 2022; Yu et al. 2022; Zhou et al. 2021) |
| | | | **Ilomastat** | | **Ilomastat** Leukaemia, Colon cancer, cancer-associated oedema (Wang et al. 2022; Zuo et al. 2022; Bissinger et al. 2021); Ohta et al. 2022) |
| | | | **ADAMTS4: US9206139, 5** | | **US9206139, 5** Rat study Osteoarthritis (Zhao et al. 2022) |

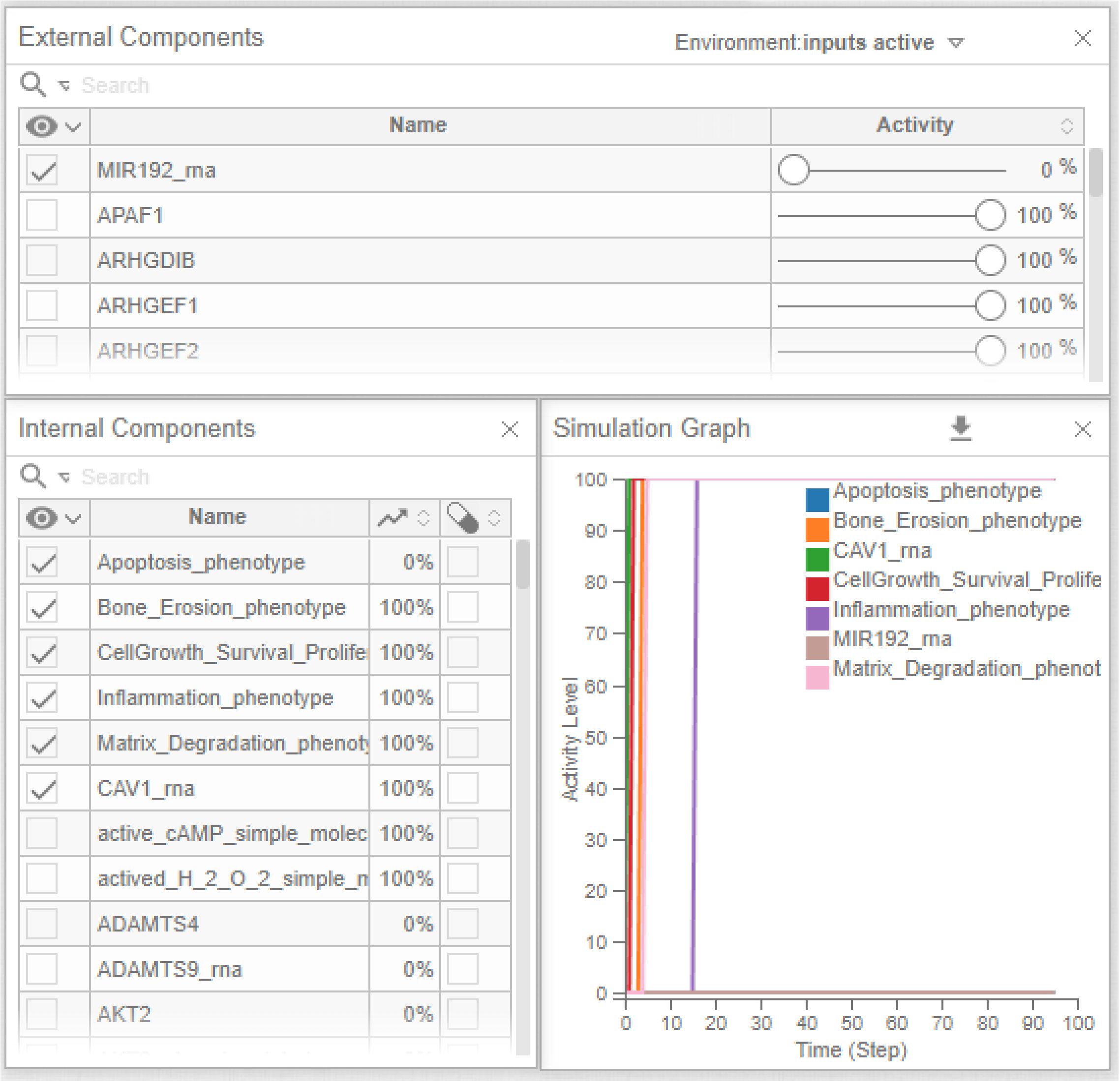


**Supplementary Figure 4**: Simulation results using the Cell Collective platform with the modified RA-FLS model. All inputs are kept active, and MIR192 is kept inactive. MIR192 is the inhibitor of CAV1, so when MIR192 is OFF, CAV1, a direct inhibitor of apoptosis, is activated. Under these conditions, apoptosis is OFF while all other phenotypes are ON, similar to the RA pathogenic state.

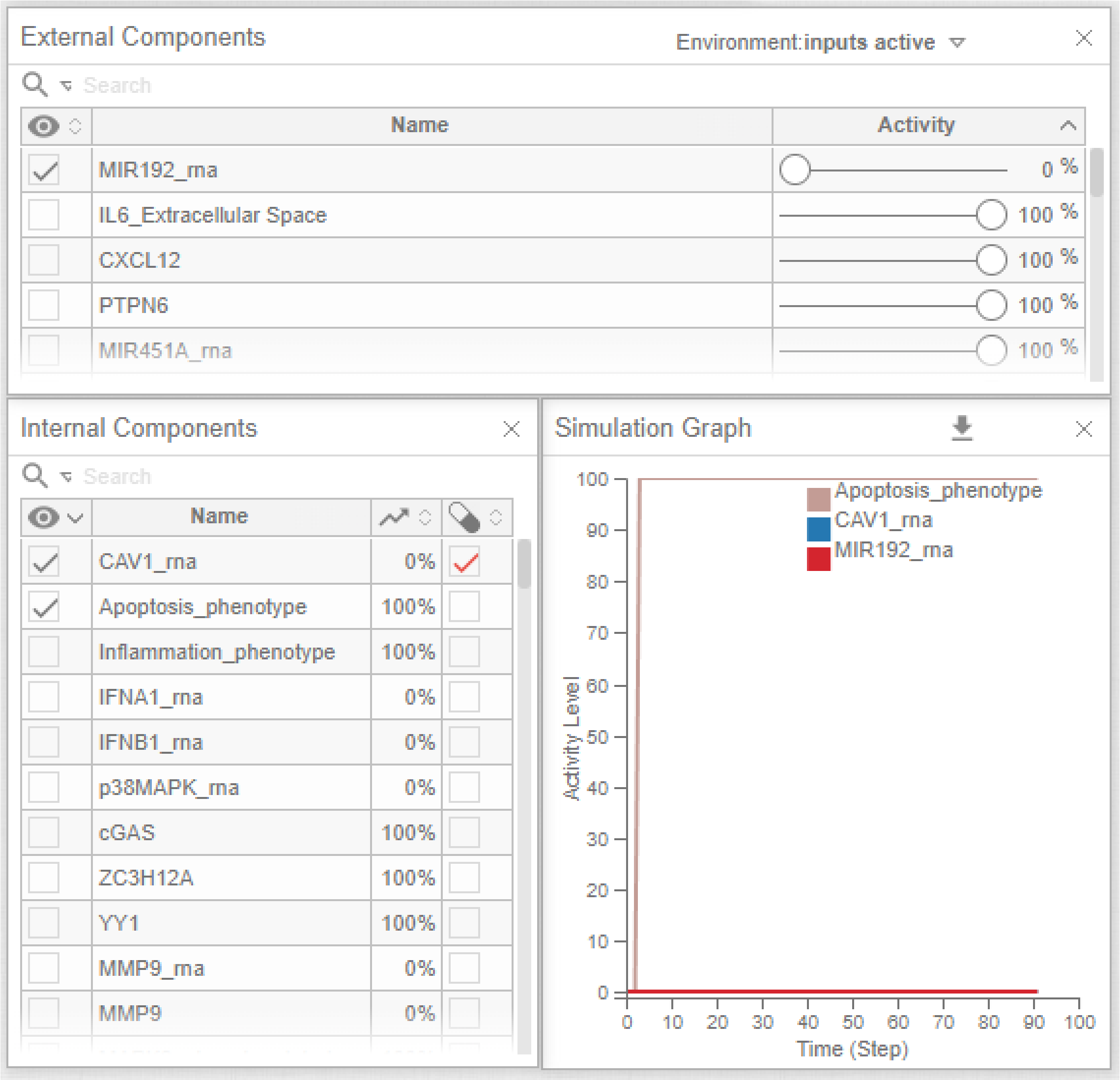


**Supplementary Figure 5**: Simulation results using the Cell Collective platform with the modified RA-FLS model. All inputs are kept active, and MIR192 is kept inactive. CAV1 is inhibited by Zoledronic acid (ZA) which results in the activation of apoptosis.

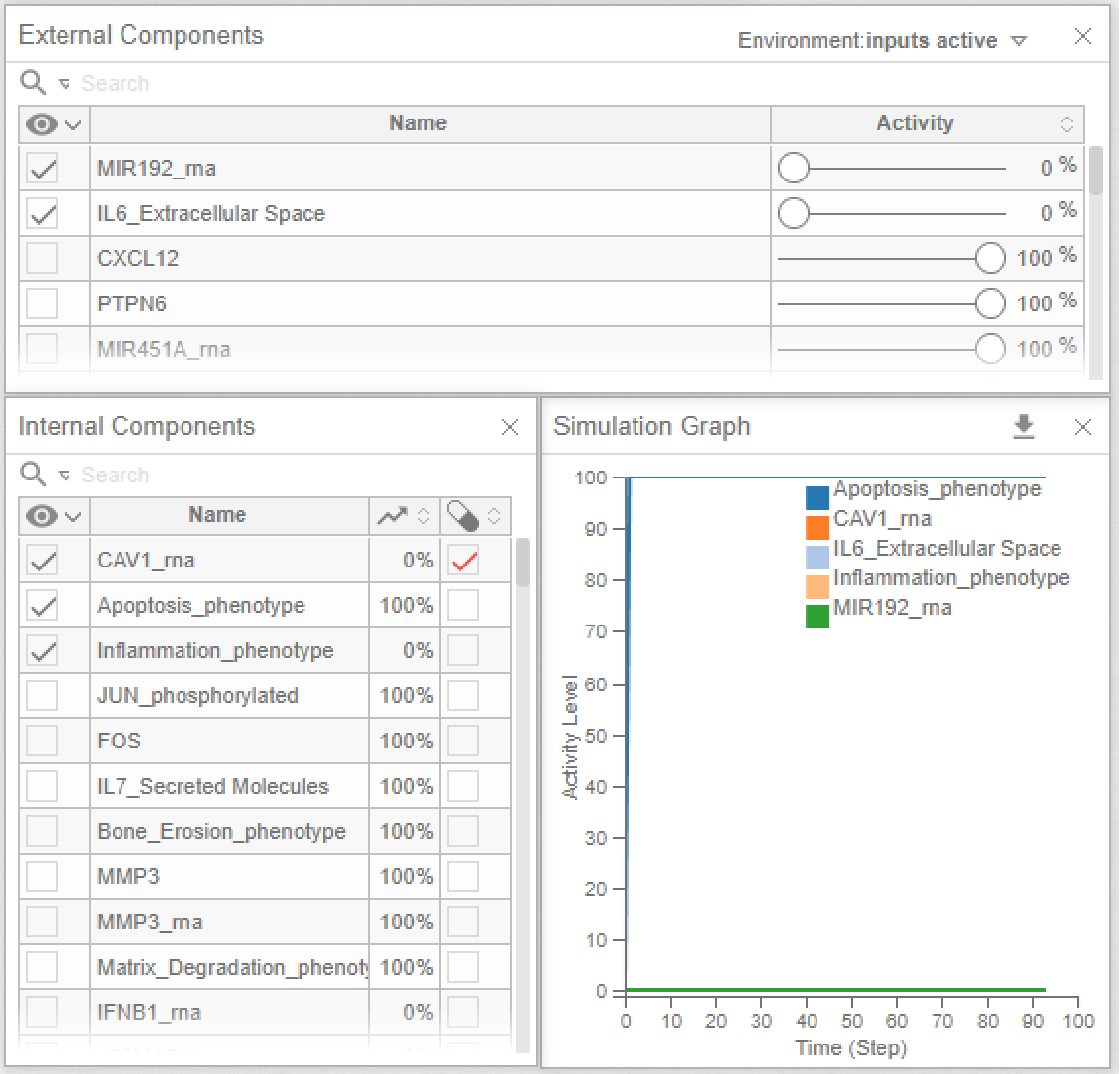


**Supplementary Figure 6**: Simulation results using the Cell Collective platform with the modified RA-FLS model. All inputs are kept active, and MIR192 is kept inactive. CAV1 is targeted by Zoledronic acid (ZA), and IL6 is targeted by Sarilumab, which results in the activation of apoptosis and the inhibition of inflammation.

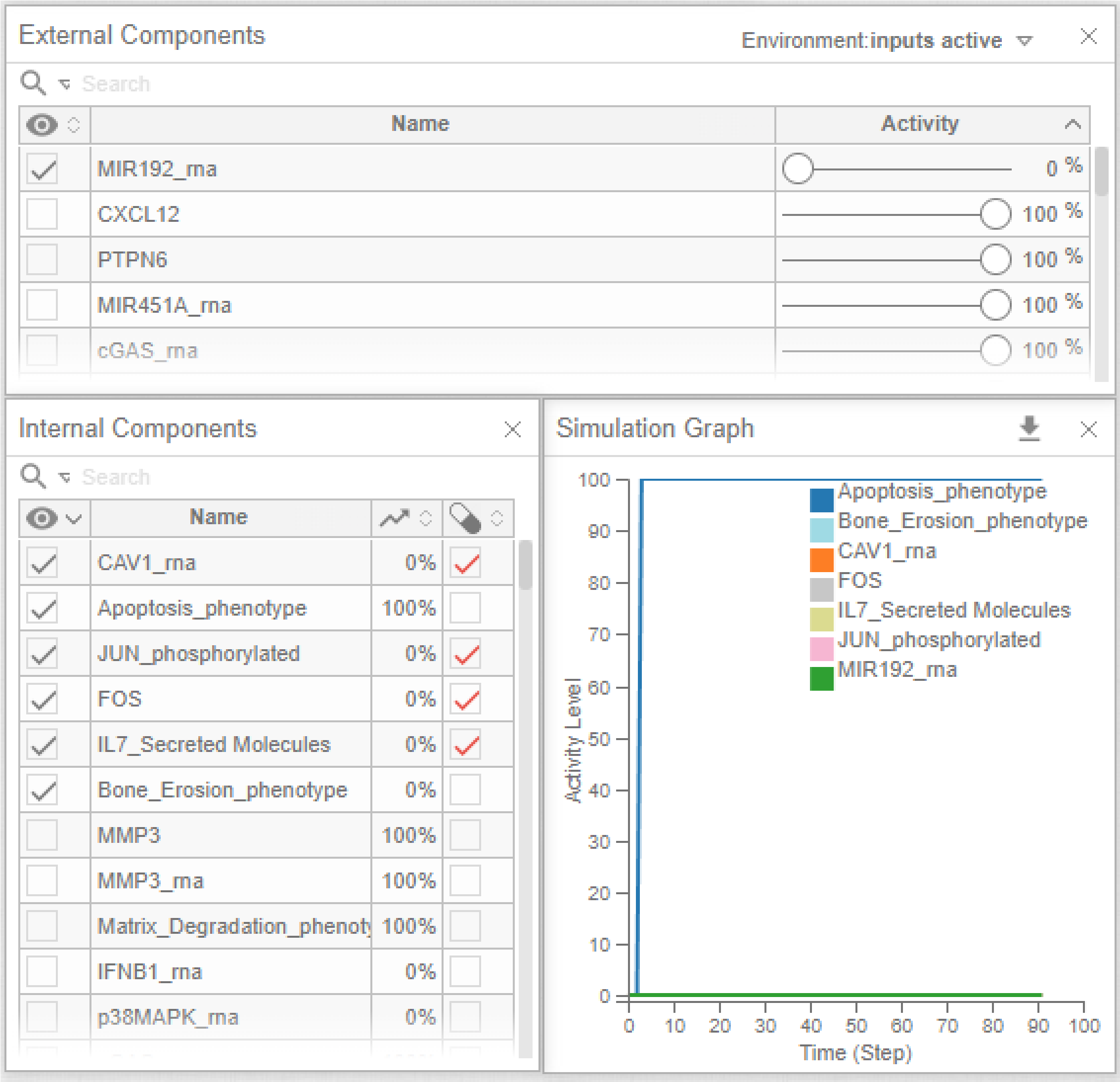


**Supplementary Figure 7**: Simulation results using the Cell Collective platform with the modified RA-FLS model. All inputs are kept active, and MIR192 is kept inactive. In addition, CAV1 is targeted by ZA, AP1 (Jun & Fos) is targeted by T-5224, and IL7 by GSK2618960/Certolizumab pegol, which results in the activation of apoptosis and the inhibition of bone erosion.

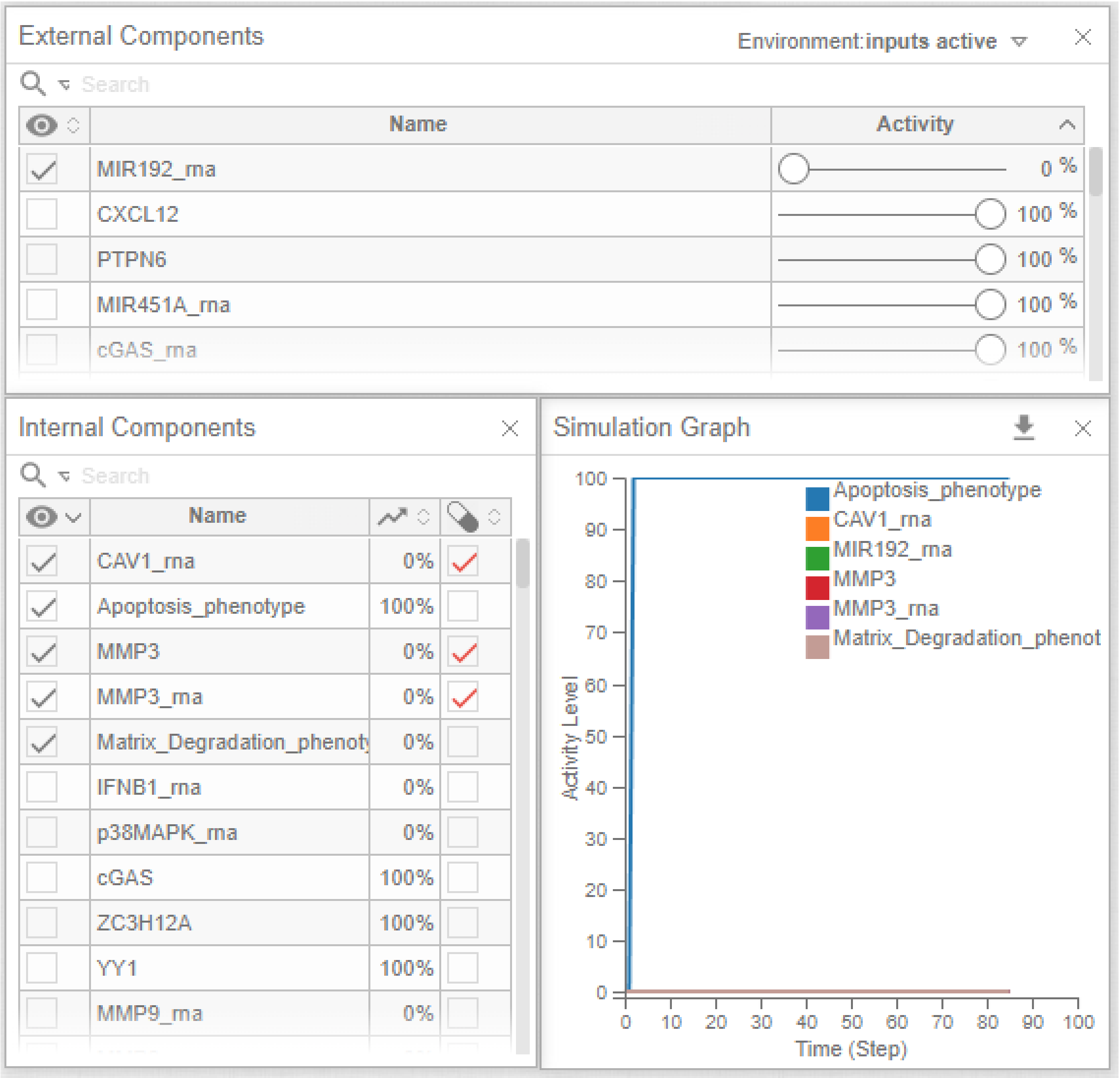


**Supplementary Figure 8**: Simulation results using the Cell Collective platform with the modified RA-FLS model. All inputs are kept active, and MIR192 is kept inactive. CAV1 is targeted by ZA, and MMP3 is targeted by Batimastat, which results in the activation of apoptosis and the inhibition of matrix degradation.

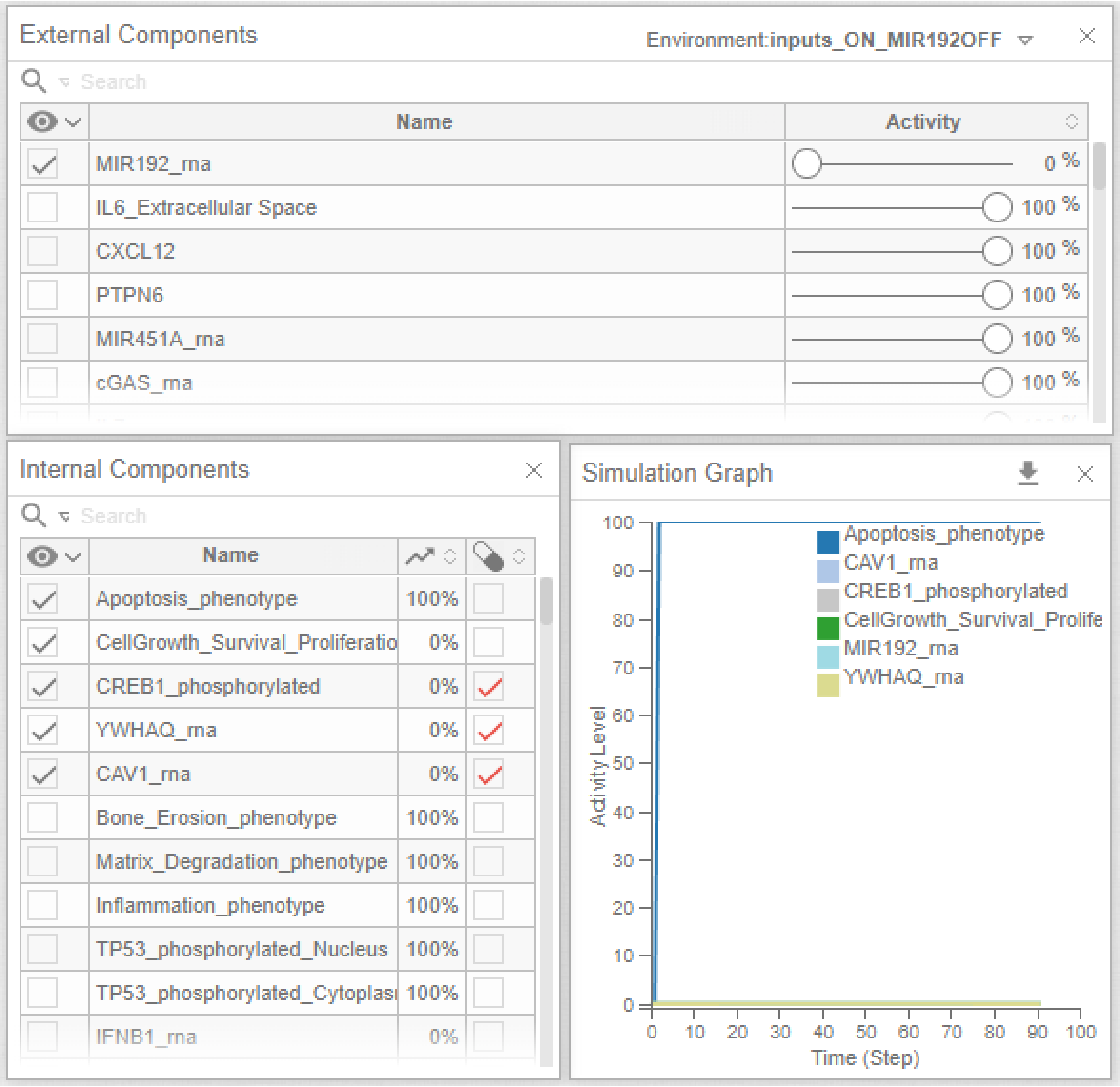


**Supplementary Figure 9**: Simulation results using the Cell Collective platform with the modified RA-FLS model. All inputs are kept active, and MIR192 is kept inactive. CAV1 is targeted by ZA, CREB1 is targeted by 666-15, and YWHAQ is targeted by AS1842856, which results in the activation of apoptosis and the inhibition of cell proliferation.
